# Genome-resolved and kinetic evidence for low-DO comammox–anammox synergy and acetate-stimulated nitrate reduction in IFAS biofilms

**DOI:** 10.64898/2026.08.17.744919

**Authors:** Zijun Meng, Juliet Johnston, Kaiqin Bian, Megan Bachmann, Mike Parsons, Francis Robinson, Charles Bott, Ameet Pinto

## Abstract

Mainstream anammox implementation for nitrogen removal is constrained by unstable nitrite supply and organic carbon requirements for nitrate byproduct removal. This study evaluated integrated fixed-film activated sludge (IFAS) biofilms to enhance anammox activity by coupling low-oxygen ammonium oxidation with volatile fatty acid (VFA)-driven nitrate reduction. Nanopore long-read metagenomic assembly recovered a high-quality, circular single-contig *Candidatus* Brocadia sapporoensis metagenome-assembled genome (MAG) from full-scale IFAS biofilms. This MAG encodes complete anammox metabolism, dissimilatory nitrate reduction to ammonium (DNRA) and acetate/propionate carbon transformation pathways. Metatranscriptomics showed that low dissolved oxygen (DO) upregulated *Ca.* B. sapporoensis genes involved in anammox, nitrate reduction, and carbon metabolism. Microaerobic assays established a DO level of 0.7 mg/L as optimal for sustaining near-maximal ammonium oxidation alongside anammox-driven total inorganic nitrogen (TIN) loss. Anoxic tests conducted in secondary effluent indicated that external acetate amendment promoted greater partial nitrate reduction and TIN loss than additional propionate amendment. Integrating this dissolved oxygen concentration with external acetate amendment in a two-stage microaerobic–anoxic system successfully achieved sequential ammonium oxidation, partial nitrate reduction, and anammox-mediated TIN removal. Stage-specific expression suggested *Ca.* B. sapporoensis could contribute to nitrite self-supplementation via *nxrAB*-mediated nitrate reduction. Overall, microaerobic ammonium oxidation and *Ca.* B. sapporoensis-driven partial nitrate reduction jointly sustain mainstream anammox activity. Furthermore, this study demonstrates that successful metabolic synergy depends fundamentally upon precise dissolved oxygen control and effective external acetate amendment.

## 1. Introduction

Anaerobic ammonium oxidation (anammox)-based wastewater nitrogen removal requires less aeration energy and organic carbon and produces less sludge than conventional nitrification–denitrification processes (Kartal et al., 2010). However, the implementation of mainstream anammox remains challenging. Ammonia oxidizing bacteria (AOB)-based partial nitritation/anammox is difficult to maintain in mainstream wastewater because low influent ammonium concentrations and competition from nitrite-oxidizing bacteria can destabilize nitrite accumulation (Y. Cao et al., 2017; Meng et al., 2025). Partial denitrification/anammox can provide nitrite for anammox bacteria through nitrate reduction, but its dependence on organic carbon addition challenges its economic and sustainability benefits (Bachmann et al., 2025).

Recent studies have reported that cooperation between complete ammonia oxidizing (comammox) bacteria and anammox bacteria can support efficient nitrogen removal from mainstream wastewater in both laboratory-scale (Cui et al., 2023; Shao and Wu, 2021; Xu et al., 2023) and full-scale systems (Johnston et al., 2024; Vilardi et al., 2023), while reducing oxygen and carbon demand and lowering biotic nitrous oxide (N_2_O) production (Kits et al., 2019). Since their discovery in 2015 (Daims et al., 2015; Pinto et al., 2015; Van Kessel et al., 2015), comammox bacteria have been widely detected in wastewater treatment systems (Annavajhala et al., 2018; Roots et al., 2019; Zheng et al., 2019). Compared with conventional AOB, comammox bacteria generally exhibit a higher affinity for ammonia (Kits et al., 2017; Sakoula et al., 2021) and better adaptation to low-oxygen environments (Koch et al., 2019; Schouten et al., 2004), conditions under which anammox bacteria can also remain active (Zhang and Okabe, 2020). In addition, both comammox and anammox bacteria are slow-growing organisms that are favored in biofilm-based systems (Kits et al., 2017), including integrated fixed-film activated sludge (IFAS) systems (Vilardi et al., 2023). These physiological and ecological characteristics suggest that comammox and anammox bacteria may occupy compatible niches under low dissolved oxygen (DO) and low-ammonium mainstream conditions. Comammox bacteria may further support anammox activity by supplying nitrite through partial nitritation under microaerobic conditions (Shao et al., 2024; Shao and Wu, 2021) or through model-predicted nitrate-reducing *Nitrospira* activity under oxygen-limited or anaerobic conditions (Martinez-Rabert et al., 2023), with nitrate originating from complete nitrification or anammox metabolism.

Anammox bacteria were traditionally considered obligate autotrophs that use CO_2_ as their sole carbon source and nitrite as the electron acceptor for converting ammonium to N_2_ (Strous et al., 1998). However, increasing evidence shows that some anammox bacteria are mixotrophic and couple volatile fatty acid (VFA) oxidation to nitrate reduction through dissimilatory nitrate reduction to ammonium (DNRA) pathway, with nitrite serving as an intermediate that can subsequently fuel anammox activity (Castro-Barros et al., 2017; Kartal et al., 2007a; Shu et al., 2015; W. Wang et al., 2022; M.-K. Winkler et al., 2012). Previous studies have shown that organic carbon addition can selectively favor the enrichment of certain anammox populations (Huang et al., 2014; Liang et al., 2015). This selective enrichment may support anammox establishment under organotrophic conditions, while VFA oxidation coupled to nitrate reduction may also contribute to influent organic carbon removal (Yin et al., 2021). In addition, previous studies suggest that acetate and propionate as prevalent VFA components in wastewater and may be preferentially utilized by distinct mixotrophic anammox populations as electron donors to drive DNRA (Chen et al., 2023; Shu et al., 2015). Low C/N ratios (i.e., 0.1–2) and the presence of ammonium can also provide anammox bacteria with a competitive advantage over heterotrophic denitrifiers and favor partial nitrate reduction to nitrite coupled with anammox, rather than complete heterotrophic denitrification or DNRA (Castro-Barros et al., 2017; Chen et al., 2016; Qiao et al., 2025; W. Wang et al., 2022).

Together, these findings suggest that comammox-anammox cooperation could enable complete nitrogen removal in mainstream wastewater systems by coupling low-DO ammonium oxidation with nitrate-reducing anammox activity. Here, nitrite required by anammox may be supplied not only from ammonia oxidation under microaerobic conditions, defined here as low liquid-phase DO below 1 mg/L (Buakaew and Ratanatamskul, 2023), but also from anoxic VFA-driven nitrate reduction, allowing residual nitrate produced by comammox, nitrite-oxidizing bacteria (NOB), or anammox metabolism to be further removed. However, the operational conditions and microbial mechanisms underlying this potential synergy remain insufficiently understood. In particular, the optimal DO concentration that balances ammonium oxidation, nitrite availability, and anammox activity under microaerobic conditions has not been systematically evaluated. Moreover, although previous studies have reported that some anammox species or genera have the potential to utilize VFAs for nitrate reduction (Castro-Barros et al., 2017), the contribution and activity of specific anammox populations in DNRA-related reactions, especially partial DNRA-anammox, across VFA types and concentrations remain unclear.

In this study, we used short- and long-read metagenomic sequencing to characterize full-scale IFAS biofilms and recovered a high-quality, circular single-contig anammox metagenome-assembled genome (MAG), identified as *Candidatus* Brocadia sapporoensis using Nanopore long-read assembly. We linked its nitrogen and carbon metabolic potential to genome-centric metatranscriptomic activity under different DO conditions. We then conducted intrinsic kinetic batch tests to identify the microaerobic DO condition most favorable for ammonium oxidation and anammox-associated nitrogen loss, and to compare the effects of external acetate and propionate amendment on anoxic nitrate-reducing/anammox-associated activity. Finally, we conducted two-stage microaerobic–anoxic batch tests by integrating the selected microaerobic condition with external acetate amendment during the subsequent anoxic stage to evaluate sequential nitrogen removal and resolve the relative contributions of comammox, AOB, anammox bacteria, and denitrifiers via metatranscriptomic analyses. These results provide genome-resolved, kinetic, and transcriptomic evidence for developing energy-efficient and potentially low-N_2_O-emission mainstream nitrogen removal processes based on comammox–anammox synergy.

## 2. Materials and Methods

### 2.1 Overview of the full-scale IFAS system

A detailed overview of the process flow diagram for the nitrogen removal process, operational parameters, and conditions is reported in previous studies (Johnston et al., 2024; Vilardi et al., 2023). However, the facility has since transitioned from an A2O configuration to a 5-stage Bardenpho configuration by incorporating a second anoxic partial denitrification-anammox (PdNA) IFAS zone within the original treatment trains (**Figure 1**). In addition, the plastic carrier media in the aerobic zone (R4) consists of a blend of approximately 90% AnoxKaldnes K3 (specific surface area 500 m^2^ m^−3^) and 10% WWW-02 media (specific surface area 800 m^2^ m^−3^) originating from the PdNA IFAS second anoxic zone (R5). The PdNA IFAS second anoxic zone (R5) now only contains WWW-01 media (specific surface area 650 m^2^ m^−3^). The aerobic IFAS zone and the PdNA IFAS second anoxic zone were operated at a carrier fill fraction of 60% and 45%, respectively.

**Figure 1.**
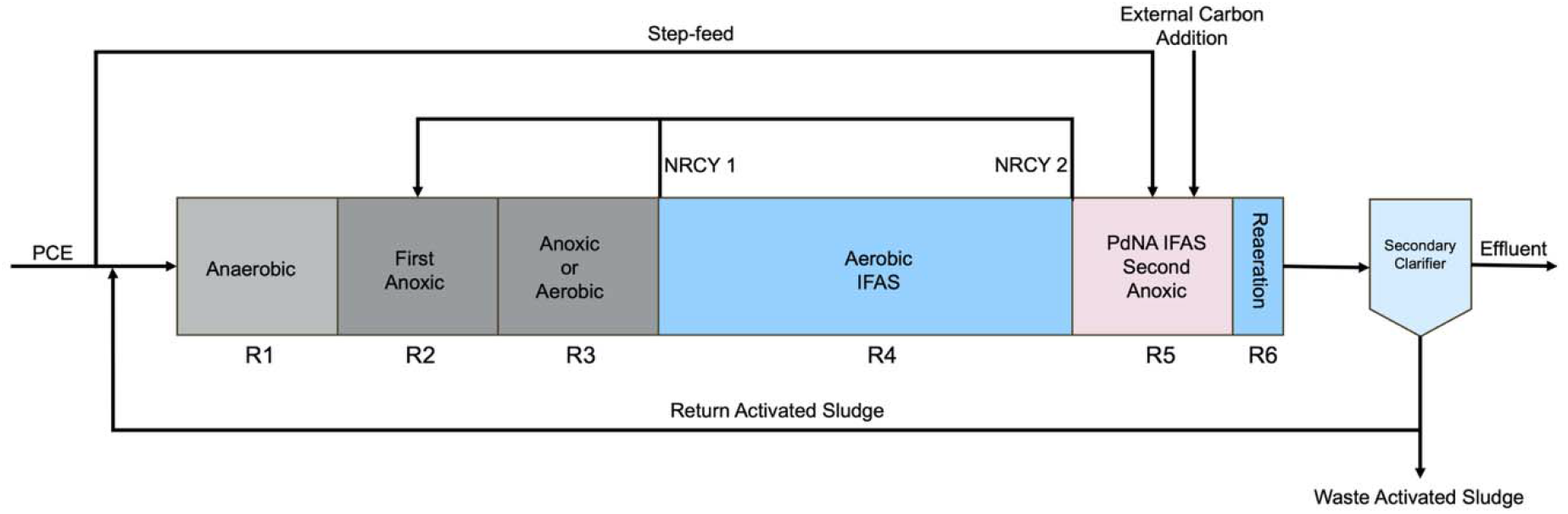
Secondary treatment configuration of full-scale IFAS system. NRCY = nitrate recycle. PCE = primary clarifier effluent.

### 2.2 IFAS media collection and batch assay

IFAS carrier media were collected from the aerobic IFAS zone of R4, transported submerged in secondary clarifier effluent, and shipped overnight to Georgia Institute of Technology. The collected media consisted of the mixed carrier population present in R4, including both K3 and WWW-02 carriers. Before the batch assays, IFAS media were washed twice with secondary clarifier effluent to remove loosely associated biomass. Separate carrier pieces were randomly selected for each batch flask, and carrier pieces were not reused between assays. Because carriers were randomly selected from the mixed R4 carrier media, the exact K3-to-WWW-02 ratio in each batch bottle or flask was not controlled. When assays were not being conducted overnight, carrier media were kept submerged in secondary clarifier effluent in the original shipping bottles and stored at 4 °C. Before each assay, the media were brought to room temperature and equilibrated with secondary clarifier effluent under mixing for at least 1 h before substrate addition.

All batch assays were conducted using secondary clarifier effluent as the liquid matrix, with continuous mixing provided by magnetic stir bars or internal magnetic stir paddles. Aqueous samples were collected immediately after substrate addition and at designated time intervals, filtered through 0.22 μm syringe filters (TISCH, Cat. No. SF14499) and stored for subsequent nitrogen species and soluble chemical oxygen demand (COD) analyses. Specific rates were normalized to attached total solids (TS) to enable comparison across biomass concentrations. Biological replicate numbers for each batch condition are summarized in **Table 1**.

**Table 1.**
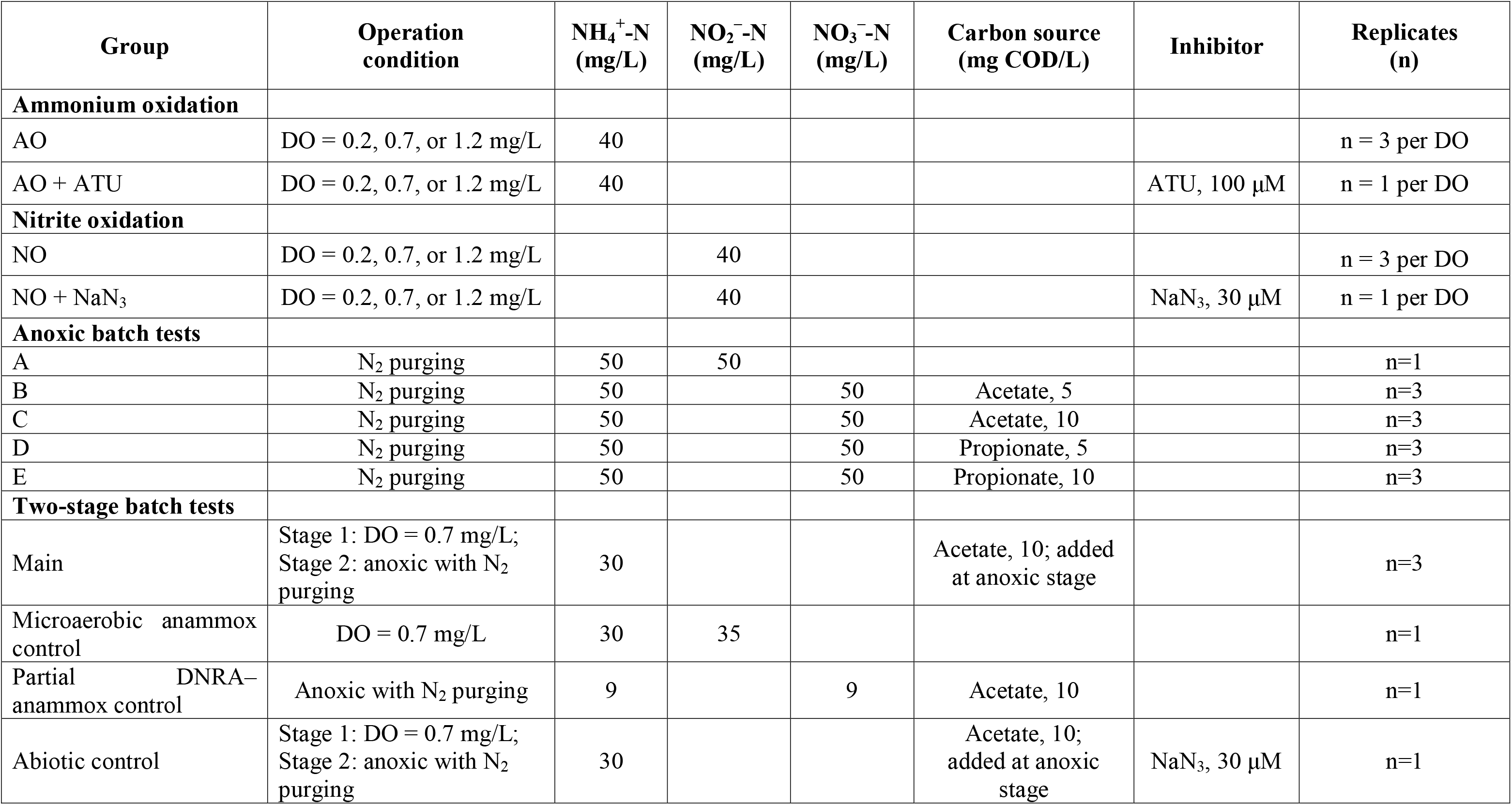
Substrate combinations, operational conditions, inhibitors, and biological replicate numbers for aerobic, anoxic, and two-stage batch assays.

| Group | Operation condition | NH<sub>4</sub><sup>+</sup>-N (mg/L) | NO<sub>2</sub><sup>-</sup>-N (mg/L) | NO<sub>3</sub><sup>-</sup>-N (mg/L) | Carbon source (mg COD/L) | Inhibitor | Replicates (n) |
| --- | --- | --- | --- | --- | --- | --- | --- |
| **Ammonium oxidation** |  |  |  |  |  |  |  |
| AO | DO = 0.2, 0.7, or 1.2 mg/L | 40 |  |  |  |  | n = 3 per DO |
| AO + ATU | DO = 0.2, 0.7, or 1.2 mg/L | 40 |  |  |  | ATU, 100 µM | n = 1 per DO |
| **Nitrite oxidation** |  |  |  |  |  |  |  |
| NO | DO = 0.2, 0.7, or 1.2 mg/L |  | 40 |  |  |  | n = 3 per DO |
| NO + NaN<sub>3</sub> | DO = 0.2, 0.7, or 1.2 mg/L |  | 40 |  |  | NaN<sub>3</sub>, 30 µM | n = 1 per DO |
| **Anoxic batch tests** |  |  |  |  |  |  |  |
| A | N<sub>2</sub> purging | 50 | 50 |  |  |  | n=1 |
| B | N<sub>2</sub> purging | 50 |  | 50 | Acetate, 5 |  | n=3 |
| C | N<sub>2</sub> purging | 50 |  | 50 | Acetate, 10 |  | n=3 |
| D | N<sub>2</sub> purging | 50 |  | 50 | Propionate, 5 |  | n=3 |
| E | N<sub>2</sub> purging | 50 |  | 50 | Propionate, 10 |  | n=3 |
| **Two-stage batch tests** |  |  |  |  |  |  |  |
| Main | Stage 1: DO = 0.7 mg/L;<br>Stage 2: anoxic with N<sub>2</sub> purging | 30 |  |  | Acetate, 10; added at anoxic stage |  | n=3 |
| Microaerobic anammox control | DO = 0.7 mg/L | 30 | 35 |  |  |  | n=1 |
| Partial DNRA–anammox control | Anoxic with N<sub>2</sub> purging | 9 |  | 9 | Acetate, 10 |  | n=1 |
| Abiotic control | Stage 1: DO = 0.7 mg/L;<br>Stage 2: anoxic with N<sub>2</sub> purging | 30 |  |  | Acetate, 10; added at anoxic stage | NaN<sub>3</sub>, 30 µM | n=1 |

### 2.3 Intrinsic kinetic batch assays and two-stage batch assays

The substrate combinations, operational conditions, inhibitors, and biological replicate numbers for all batch assays are summarized in **Table 1**.

Aerobic intrinsic kinetic assays were conducted in 1.3 L Corning disposable spinner flasks (Cat. No. 3580) containing 40 IFAS carrier pieces and 1.3 L secondary clarifier effluent. Three target bulk DO setpoints were tested: 0.2, 0.7, and 1.2 mg/L. DO concentrations were controlled by regulating aeration from a pump connected to a flowmeter (COKYIS, 0.1–1.5 L/min) and simultaneous N_2_ purging. DO was monitored in real time using a Thermo Scientific Orion Star A329 portable meter (Cat. No. STARA3290), and air and N_2_ flow rates were adjusted to maintain the target bulk DO concentrations. Differential inhibition controls were conducted to support the interpretation of aerobic ammonia and nitrite oxidation pathways. Allylthiourea (ATU; 100 μM) was added 30 min before ammonium addition to inhibit aerobic ammonia oxidation by AOB and comammox bacteria (Ali, 2013; Ginestet et al., 1998), and sodium azide (30 μM) was added 30 min before nitrite addition to inhibit aerobic nitrite oxidation by NOB and comammox bacteria (Ginestet et al., 1998). Samples were collected immediately after substrate addition and every 20 min over the 2 h reaction period.

Anoxic intrinsic kinetic assays were conducted in 1 L Pyrex media bottles containing 30 IFAS carrier pieces and 900 mL secondary clarifier effluent. Bottles were fitted with two-port caps, continuously purged with N_2_ to maintain anoxic conditions, and mixed throughout the assay period using magnetic stir bars. Reactors were purged with N_2_ for 30 min before substrate addition and continuously flushed with N_2_ throughout the 3 h reaction period. Samples were collected immediately after substrate addition and every 30 min over the 3 h reaction period. pH and temperature were measured before substrate addition and at the end of each assay using a Thermo Scientific Orion Star A329 portable meter.

Two-stage microaerobic–anoxic batch tests were conducted in 1.3 L Corning disposable spinner flasks containing 40 IFAS carrier pieces and 1.3 L secondary clarifier effluent. The reactor was first operated at a target bulk DO of 0.7 mg/L for 2 h with ammonium as the sole added nitrogen source, which is referred to here as the microaerobic stage. Aeration was then stopped, N_2_ purging was initiated for 20 min to establish anoxic conditions, and acetate was added as the external carbon source. The reactor was subsequently operated under anoxic conditions for an additional 2 h. Three additional control assays were included: a microaerobic anammox control, an anoxic partial DNRA–anammox control, and an abiotic control with sodium azide added at the start. Samples were collected immediately after initial substrate addition and every 20 min during both stages for nitrogen species and soluble COD analyses. At the transition between the microaerobic and anoxic stages, samples were collected both before and immediately after acetate addition to determine the pre-acetate baseline and the initial acetate-amended COD concentration

### 2.4 Water quality analysis and calculations

Ammonium concentrations in filtered aqueous samples were measured according to Standard Methods (American Public Health Association and others, 2005). Nitrite and nitrate concentrations were quantified using a Metrohm 930 Compact IC Flex ion chromatograph (Metrohm, Herisau, Switzerland), and soluble COD was measured using low-range HACH COD digestion vials (HACH, Loveland, USA). Attached biomass on IFAS media was quantified as total solids (TS). Briefly, media pieces were dried at 105 °C, weighed, scrubbed with 2 N H_2_SO_4_ to remove attached biomass, dried again at 105 °C, and reweighed. Attached TS was calculated as the difference between dried media with biomass and cleaned media, and the total attached biomass in each assay was estimated from the average TS per media piece and the number of media pieces used. Total inorganic nitrogen (TIN) was calculated as the sum of NH_4_^+^–N, NO_2_^−^–N, and NO_3_^−^–N at each time point. Temporal changes in nitrogen species, TIN, and soluble COD were used to calculate volumetric transformation rates. Specific rates were calculated by normalizing volumetric rates to the attached biomass concentration and are reported as mg N/g TS-h or mg COD/g TS-h.

### 2.5 Nucleic Acid Extraction

For DNA extraction, carrier media were collected only from the main triplicate two-stage batch tests at the end of the anoxic stage. Because the same batch of IFAS media was used for these assays, three to five carrier pieces were collected from each replicate at the end of the complete two-stage operation and transferred into sterile 50 mL centrifuge tubes. Samples were stored at −80 °C until DNA extraction. At the end of each batch experiment, three carrier media pieces were collected and transferred into sterile 50 mL centrifuge tubes and stored at −80 °C specifically for DNA extraction. For DNA extraction, five pieces of media from each DNA sample were immersed in 1× phosphate-buffered saline (PBS) for 30 min to remove non-adherent bacteria. The carriers were then transferred into a 50 mL sterile centrifuge tube containing 25 mL of sterile PBS (VWR PBS Powder) and vortexed vigorously for 10 min to detach attached biomass and biofilm from the media surface. The resulting suspension was concentrated by centrifugation at 6000 rpm for 14 min, and the pellet was collected into PCR-grade Eppendorf tubes. Approximately 50 mg of biomass from each sample was used for DNA extraction using the Qiagen DNeasy PowerSoil Pro Kit (Cat. No. 47016) on a Qiacube system (Cat. No. 9002160). Extracted DNA was quantified using a Qubit 4 fluorometer following standard high-sensitivity dsDNA protocols (ThermoFisher Scientific, Waltham, Massachusetts, USA).

For RNA-based analyses, IFAS carrier media were collected separately from DNA samples. In the main two-stage batch tests, three to five carrier pieces were collected at the beginning of the microaerobic stage, at the end of the microaerobic stage before acetate addition, and at the end of the anoxic stage. For the microaerobic anammox control and the anoxic partial DNRA–anammox control, three to five carrier pieces were collected at the beginning and end of each 2 h assay. Immediately after collection, each RNA sample was submerged in DNA/RNA Shield (Zymo, Irvine, California, USA) and stored at −80 °C until RNA extraction. RNA was extracted using protocols from the Quick-RNA fecal/soil microbe microprep kit with some modifications (Zymo, Irvine, California, USA) as described previously (Johnston et al., 2024).

### 2.6 Metagenomic Sequencing and Data Processing

Two DNA samples extracted from the main two-stage batch tests were submitted to the Georgia Institute of Technology Molecular Evolution Core for short-read sequencing on the Element Bioscience AVITI platform. The same DNA samples were also sequenced on the Oxford Nanopore MinION Mk1C platform in our laboratory. Library preparation and sequencing details for both short-read and long-read sequencing are provided in **Text S1**. Raw short reads were filtered using fastp v0.24.1 (Chen, 2023) and the Univec database was used to remove contamination from the filtered reads. Raw Nanopore reads were visualized using Nanoplot v1.46.2 (De Coster and Rademakers, 2023) and then demultiplexed, with barcode and adapter sequences trimmed using qcat v1.1.0 (https://github.com/nanoporetech/qcat). Following demultiplexing, the Nanopore reads were further processed using Chopper v0.9.2 (De Coster and Rademakers, 2023), to remove reads shorter than 1000 bp and reads with an average Phred quality score below 12. Long-read-only assemblies were generated using metaFlye v2.9.5 (Kolmogorov et al., 2020), followed by removal of contigs shorter than 4000 bp using seqtk v1.4-r122 (https://github.com/lh3/seqtk) and polishing with Medaka v2.0.1 (https://github.com/nanoporetech/medaka) using the r1041_e82_400bps_sup_v5.0.0 model. Medaka-polished assemblies were further polished using cleaned short reads with Polypolish v0.6.1 (Wick and Holt, 2022) and Pypolca v0.4.0 (Bouras et al., 2024; Zimin and Salzberg, 2020). Both Medaka-polished long-read assemblies and short-read-polished assemblies were used for genome binning and subsequent quality assessment.

Genome binning was performed using MetaBAT2 v2.17 (Kang et al., 2019), SemiBin2 v2.3.0 (Pan et al., 2022), and Vamb v5.0.4 (Nissen et al., 2021). DAStool v1.1.7 (Sieber et al., 2018) was used to integrate and dereplicate bins generated by individual binning tools to obtain a nonredundant MAG set. MAG quality was assessed using CheckM2 v1.1.0 (Chklovski et al., 2023), and taxonomy was assigned using Genome Taxonomy Database Toolkit (GTDB-Tk 2.6.1, database release R226) (Chaumeil et al., 2020; Parks et al., 2018). Final MAGs were selected for downstream analyses based on contiguity, completeness, and contamination across bins recovered from both Medaka-polished long-read assemblies and short-read-polished assemblies. Subsequently, genes were predicted and annotated using Bakta v1.12 (Schwengers et al., 2021), and functional annotation against KEGG orthology was performed using EnrichM v0.5.0 (https://github.com/geronimp/enrichM). The EnrichM database incorporated a KO-annotated UniRef100 database and KoFamKOALA Hidden Markov Models for functional annotation. The EnrichM “classify” function was used to identify complete nitrogen-cycling KEGG Orthology modules and key nitrogen and carbon metabolism genes present in the MAGs. Phylogenomic reconstruction was performed for medium- and high-quality MAGs recovered from the IFAS biofilm communities. A maximum-likelihood phylogenetic tree was inferred using IQ-TREE v2.4.0 (Minh et al., 2020) with the LG + R5 model and 1,000 bootstrap replicates, and the tree was visualized using TVBOT v2.6.1 (Xie et al., 2023). The breadth of coverage and relative abundance of each MAG were calculated using CoverM v0.7.0 (https://github.com/wwood/CoverM) based on short-read mapping. MAG relative abundance was calculated from reads per kilobase per million mapped reads (RPKM), and percent relative abundance was calculated by dividing reads mapped to each MAG by the total reads in each sample.

### 2.7 Metatranscriptomic Sequencing and Data Processing

Raw metatranscriptomic RNA sequencing reads generated in our previous study across different DO operational conditions and reactor locations (R1, R3, and R4) were reused in this study (Johnston et al., 2024). Sequencing protocols and data generation details are described in the original publication. In addition, RNA samples collected from the two-stage aerobic–anoxic batch tests and associated control assays were sequenced for metatranscriptomic analysis in this study. Extracted RNA samples were submitted to the Georgia Institute of Technology’s Molecular Evolution Core for rRNA depletion and RNA sequencing. RNA integrity, read length, and quantification were performed on a Tape Station 4200 (Agilent, Santa Clara, California, USA). The average RNA integrity number (RIN) value was 5.48 ± 0.41 across all samples. Excess rRNA was depleted using the QiaSeq FastSelect 5S/16S/23S depletion kit as well as the FastSelect HMR rRNA probes (Qiagen, Hilden, Germany). RNA sequencing was performed on an AVITI high output mode PE150bp instrument to obtain 976 million read pairs (Element Biosciences, San Diego, California, USA). Samples were demultiplexed, and barcodes were removed by the core facilities prior to releasing sequences.

Raw reads from both the previously generated and newly sequenced metatranscriptomic datasets were processed using the same bioinformatic workflow. Raw reads were first preprocessed using fastp v0.24.1 (Chen, 2023), followed by separation of mRNA reads from residual rRNA using SortMeRNA v4.3.7 (Kopylova et al., 2012). The remaining mRNA reads were competitively mapped using Minimap2 v2.28 (r1209) (Li, 2018) against a concatenated reference FASTA containing a nonredundant set of nitrogen-cycling MAGs. The reference set included ammonia-oxidizing bacterial, comammox, nitrite-oxidizing bacterial, and anammox MAGs; MAGs harboring complete canonical denitrification pathway; MAGs harboring complete DNRA metabolic pathway; and partial denitrification MAGs containing *narGHI* or *napAB*. Mapped reads were converted to BAM format, sorted, and indexed using SAMtools v1.21 (Danecek et al., 2021). MAGs were annotated using bakta v1.12 (Schwengers et al., 2021), and the mapped mRNA reads were quantified as gene-level read counts based on the annotated MAG features using dirseq v0.4.3 (Woodcroft et al., 2018).

### 2.8 Statistical analyses

Statistical analyses and figure generation were performed in R v4.6.0 (R Core Team, 2021). Transformation rates were estimated using linear regression of concentration changes over time. For comparisons involving more than two experimental conditions, one-way analysis of variance (ANOVA) followed by Tukey’s honest significant difference (HSD) (Keselman and Rogan, 1977) post hoc test was used to identify significant pairwise differences. For comparisons between two conditions, Welch’s two-sample t-test was used. Differential gene expression analysis was performed with DESeq2 (Love et al., 2014). Within DESeq2, log2 fold changes were used to determine the direction and magnitude of transcriptional responses, and Benjamini–Hochberg adjusted *p* values were used to determine statistical significance. Differences were considered statistically significant at an *p* or adjusted *p*-value < 0.05. For all figures and tables, significance levels are denoted as follows: *p* or *padj* < 0.05 (*), *p* or *padj* < 0.01 (**), and *p* or *padj* < 0.001 (***), depending on the statistical test reported.

## Results and Discussion

### 3.1 High-quality anammox MAG encodes DNRA-associated genes and exhibits *nxrAB* upregulation under low DO conditions

DNA extracted from IFAS carrier media biofilms collected from aerobic zone R4 were subject to Illumina and Nanopore sequencing. Following hybrid assembly and binning, 276 MAGs were recovered, of which 209 were classified as medium to high quality (≥50% completeness, ≤10% contamination) (The Genome Standards Consortium et al., 2017). Among the 209 medium- and high-quality MAGs, 181 encoded at least one gene associated with nitrogen metabolism. A subset of 88 MAGs across eight phyla were selected for phylogenomic analysis because they encoded either a complete nitrogen removal metabolic pathway or the functional potential for nitrate reduction to nitrite, defined here as partial denitrification (**Figure 2**). Only one *Nitrospira*-comammox and one *Nitrosomonas*-AOB MAGs were recovered, with low relative abundances of 0.31% and 0.40%, respectively, consistent with our previous observations from the same IFAS system (Vilardi et al., 2023). FastANI-based pairwise genome comparison showed that the recovered comammox MAG shared >95% average nucleotide identity with *Ca.* Nitrospira nitrosa and with comammox MAGs previously recovered from the same IFAS system and other wastewater treatment systems (**Figure S1**) (Cotto et al., 2023). Five MAGs spanning Pseudomonadota and Nitrospirota were identified as NOB, where four of these were classified as *Nitrospira*-NOB and together accounted for 5.36% relative abundance, representing the dominant nitrifying group in this IFAS system. In addition, fifteen MAGs spanning Pseudomonadota, Actinomycetota, Acidobacteriota, and Planctomycetota encoded complete DNRA pathways, whereas seven MAGs within Pseudomonadota harbored complete denitrification pathways. These findings are consistent with previous studies showing that members of Pseudomonadota frequently co-occur with Planctomycetota in engineered bioreactor systems (Lawson et al., 2017; Yang et al., 2018) and commonly possess genetic potential for both denitrification and DNRA metabolism (Pan et al., 2020; Zhao et al., 2019). MAGs from all seven phyla other than Nitrospirota also encoded nitrate reduction genes, indicating that the potential for nitrate reduction to nitrite was phylogenetically widespread within the IFAS community.

**Figure 2.**
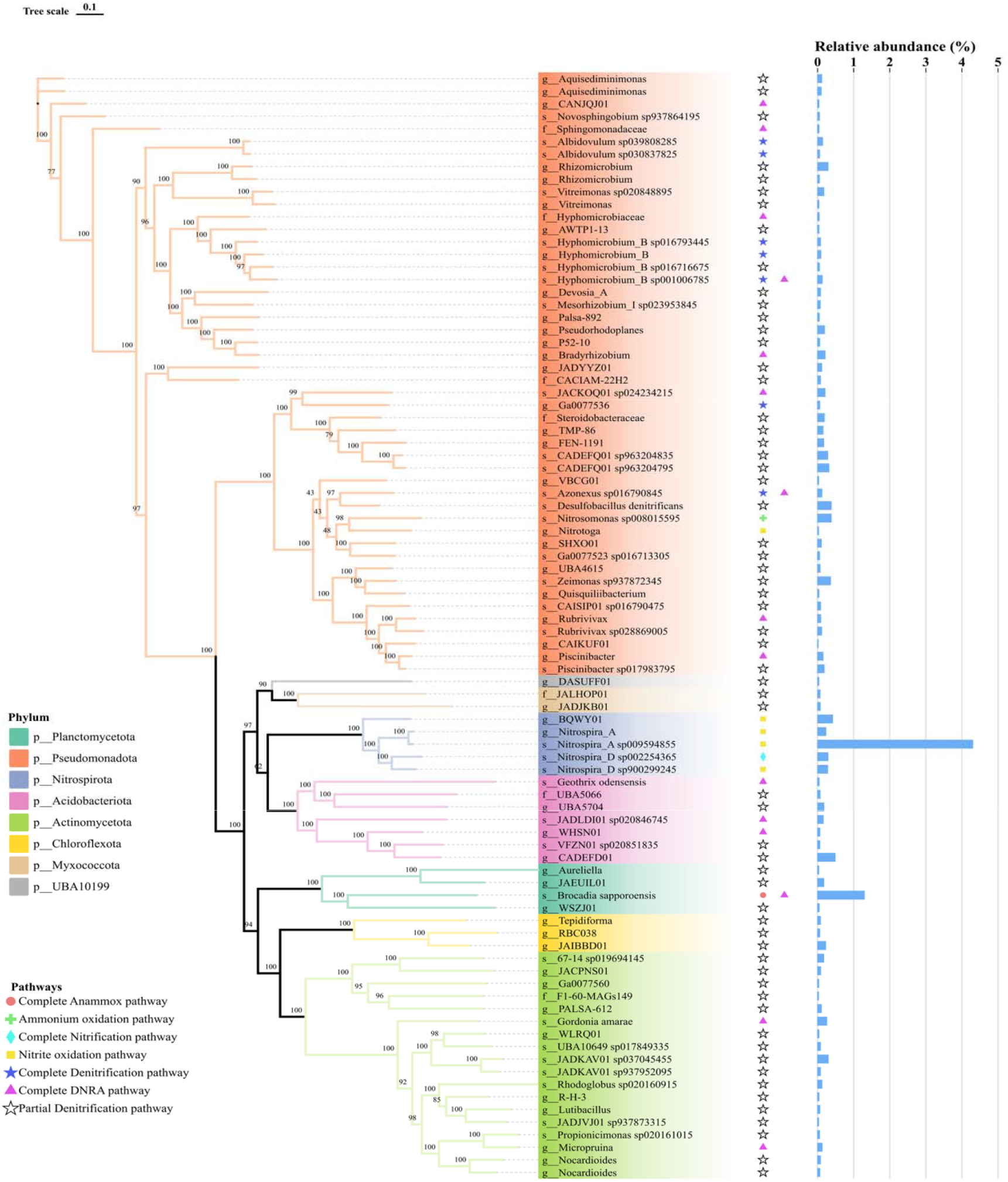
Phylogenomic reconstruction of selected MAGs with nitrogen-cycling potential recovered from aerobic IFAS biofilm. MAGs were selected for visualization based on the presence of complete nitrogen-cycling pathways or genes supporting nitrate reduction to nitrite, defined here as partial denitrification. A maximum-likelihood phylogenetic tree was inferred using IQ-TREE v2.4.0 (Minh et al., 2020) with the LG + R5 model and 1,000 bootstrap replicates. Branch colors and background shading indicate phylum-level taxonomy, symbols denote predicted nitrogen transformation pathways, and bars on the right show MAG relative abundance in the IFAS biofilm metagenome.

Notably, one of the more abundant nitrogen-cycling MAGs, with a relative abundance of 1.32%, was classified as *Ca.* Brocadia sapporoensis (Planctomycetes). This high-quality *Ca.* Brocadia sapporoensis MAG was recovered from the Nanopore long-read assembly and comprised a single circular contig of 3.49 Mb, with an estimated completeness of 100.00% and contamination of 0.58%. Circularity was supported by a raw Nanopore read spanning the terminal-to-initial contig junction and by reproducible metaFlye assembly output that labeled the corresponding contig as circular. Functional annotation revealed the presence of genes associated with anammox, nitrate reduction, and DNRA processes as the primary nitrogen metabolism pathways. Specifically, *Ca.* Brocadia sapporoensis encoded hallmark anammox genes—hydroxylamine oxidoreductase (*hao*), hydrazine synthase (*hzs*), and hydrazine oxidoreductase (*hzo*)—as well as nitrite oxidoreductase genes (*nxrAB*) and nitrite reductase genes (*nrfAH*), indicating the genomic potential for a DNRA pathway enabling the reduction of nitrate to ammonium in addition to canonical anammox metabolism (Castro-Barros et al., 2017). Previous studies have also reported that DNRA-related functional genes were observed in other anammox species, including *Ca.* Brocadia sinica (Shu et al., 2016), *Ca.* Kuenenia stuttgartiensis (Chen et al., 2023), and *Ca.* Jettenia caeni (Feng et al., 2019). In addition, *Ca.* Brocadia sapporoensis lacked *nirS* and *nirK*, nitrite reduction genes involved in conversion of nitrite to NO, and this differentiates it from other bacteria that can conduct nitrite reduction (Qiao et al., 2025).

Our previous studies indicated that the activity of most nitrogen metabolism pathways in the IFAS biofilm was significantly higher under the lower bulk DO condition of 2 mg/L than under 6 mg/L (Johnston et al., 2024; Vilardi et al., 2023). Therefore, the metatranscriptomic data obtained from various reactor zones in these previous studies were mapped to the refined MAG in this study to further quantify expression of key nitrogen metabolism-related genes under varying DO conditions. Metatranscriptomic analysis indicated that most nitrogen cycling genes from the *Ca.* Brocadia sapporoensis MAG were not differentially expressed due to DO concentration changes in suspended sludge (SS) biomass. In contrast, both *nxrA* (*Tukey p* = 1 × 10^−4^) and *nxrB* (*Tukey p* = 3.42 × 10^−2^) were significantly more expressed in IFAS biofilm as compared to the suspended sludge biomass under lower bulk DO conditions (2 mg/L). This suggests that *Ca.* Brocadia sapporoensis in the biofilm phase could rely on nitrate reduction to supplement its own nitrite requirements when competing with NOB in the aerobic zone, where increased oxygen availability may allow NOB to continuously oxidize nitrite to nitrate, reducing nitrite accumulation for anammox (Xu et al., 2023). In addition, most pathways showed significantly increased expression under low bulk DO conditions in the biofilm (*p* < 0.05): *hao*/*hzo* increased 1.2-fold, *hzsA* by 1.64-fold, *nxrA* by 1.65-fold, and *nxrB* by 1.34-fold. In contrast, expression of *nrfA* and *nrfH* remained extremely low across reactor zones or DO concentrations, indicating that full DNRA (nitrate to ammonium) is unlikely to occur under such operational conditions. In addition, the expression of *hzo*/*hao* and *hzs* remained high across DO concentrations in IFAS biomass, even when the bulk DO concentration reached 6 mg/L. This suggests that anammox transcriptional activity was maintained under such DO operational conditions, likely due to the compact biofilm structure, which may impose strong diffusion resistance and limits oxygen penetration into the biofilm matrix, thereby creating anoxic microenvironments that can accommodate and protect anammox bacteria (Z. Wang et al., 2022). Together, these results provide genomic and transcriptomic evidence that *Ca.* Brocadia sapporoensis in IFAS biofilms can reduce nitrate via a partial DNRA pathway and may indeed be supplementing its nitrite requirements. By internally generating nitrite through nitrate reduction, this anammox species may alleviate nitrite limitation in competitive environments, particularly under low DO.

### 3.2 Metagenomic and metatranscriptomic analyses suggest carbon metabolic flexibility in *Ca.* Brocadia sapporoensis

Metagenomic analysis revealed the presence of several key functional genes potentially involved in acetate and propionate transformation in *Ca*. Brocadia sapporoensis (**Figure 3A–B**). These pathways can be broadly classified into direct activation of acetate or propionate to their corresponding CoA derivatives, acyl-phosphate-mediated interconversion, and a phosphoketolase-associated branch centered on acetyl-phosphate. Direct acyl-CoA formation may provide activated carbon intermediates for energy metabolism and biosynthesis, whereas the reversible *ackA*–*pta* pathway may enable interconversion among acetate, acetyl-phosphate, and acetyl-CoA in response to substrate availability and cellular energetic demands (Dittrich et al., 2008; Krivoruchko et al., 2015). The *xfp*-associated route can cleave xylulose-5-phosphate into glyceraldehyde-3-phosphate and acetyl-phosphate (Meile et al., 2001). The former can enter glycolytic metabolism, whereas the latter can be converted by *ackA* to acetate with concomitant ATP generation or potentially directed toward acetyl-CoA through *pta*, thereby linking pentose phosphate metabolism with energy conservation and acetyl-CoA-related central metabolism (Kleijn et al., 2005; Papini et al., 2012). Propionate transformation appeared to involve analogous direct activation and propionyl-phosphate-mediated routes but lacked an identifiable *xfp*-associated route. In addition, *por* may connect acetyl-CoA and propionyl-CoA metabolism with pyruvate and 2-oxobutyrate formation, respectively, providing potential entry points into central carbon metabolism and biosynthetic pathways (Ji et al., 2021; Lawson et al., 2021). Previous studies have reported substantial variation in acetate-associated metabolic potential among anammox bacteria (Chen et al., 2023; Feng et al., 2019; Yin et al., 2021; Zhang et al., 2021). Specifically, *Ca.* Jettenia caeni (Feng et al., 2019) and *Ca.* Kuenenia stuttgartiensis (Chen et al., 2023) were reported to encode an *acs* pathway and an aldehyde dehydrogenase (*ALDH*, EC 1.2.1.3)-associated route catalyzing the reversible conversion between acetate and acetaldehyde. Pathway composition also varied within the genus *Brocadia*: the *Ca.* Brocadia sapporoensis MAG recovered here encoded *acs*, *ackA*–*xfp*, and *ackA*–*pta* routes, whereas *Ca.* Brocadia sinica was reported to possess *acs*, *ALDH*, and *ackA*–*xfp* routes (Feng et al., 2019), and *Ca. Brocadia fulgida* encoded only *acs* and *ackA*–*pta* pathways (Yin et al., 2021). Together, these differences indicate species-level variation in the metabolic potential for acetate transformation among anammox bacteria.

**Figure 3.**
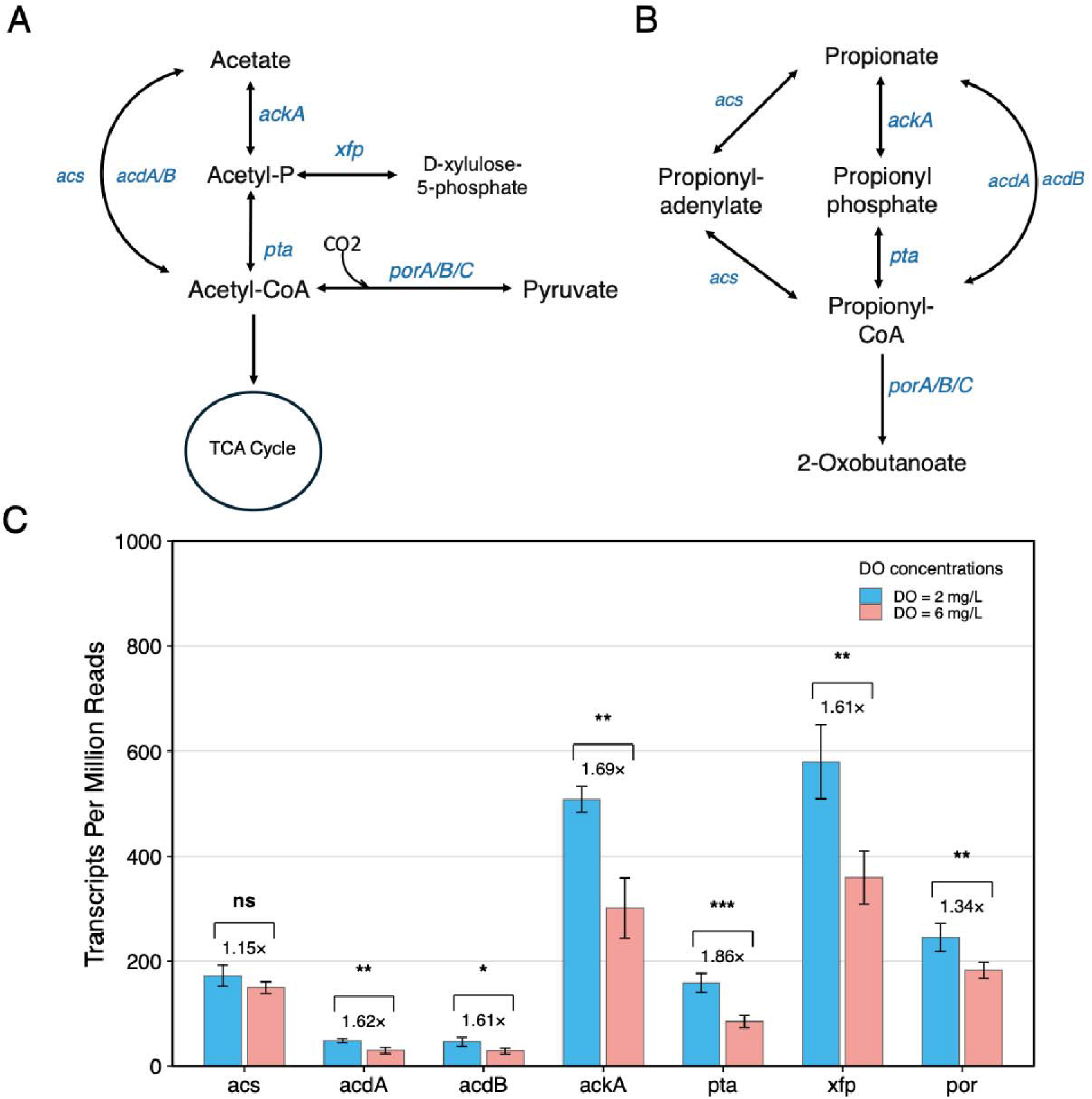
Genome-inferred acetate-associated (A) and propionate-associated (B) metabolic pathways, and (C) transcriptional expression of key functional genes in *Ca.* Brocadia sapporoensis within IFAS biofilm under different DO conditions in R4 aerobic zone.

Metatranscriptomic analysis further indicated that the carbon-associated pathways were transcriptionally active in *Ca.* Brocadia sapporoensis within the IFAS biofilm (**Figure 3C**). Among the examined genes, *ackA* and *xfp* exhibited relatively high transcripts per million (TPM) values under both DO conditions, indicating that functions associated with the acetyl-phosphate node were transcriptionally prominent in *Ca.* Brocadia sapporoensis. In addition, expression of AMP-forming *acs* was markedly higher than that of ADP-forming *acdA* and *acdB*. Both enzymes can catalyze acetate into acetyl-CoA, but ADP-forming *acdA* and *acdB* catalyzes the synthesis of acetyl-CoA from acetate in a single step, while AMP-forming *acs* synthesizes it in two steps (Starai and Escalante-Semerena, 2004), and such different mechanisms between these two groups of enzymes in *Ca.* Brocadia sapporoensis may be one reason for their differing expression levels. Though *pta* showed relatively lower expression compared to *xfp*, its detectable expression provided transcriptional support for mediating interconversion between acetyl-phosphate and acetyl-CoA. The relatively high TPM values of *por* further suggest that the generated acetyl-CoA and propionyl-CoA may be further transformed to pyruvate and 2-oxobutanoate, respectively, thereby providing potential intermediates for central carbon metabolism and amino acid biosynthesis (Ji et al., 2021; Ragsdale, 2003). Notably, most of the carbon-associated genes showed significantly increased expression under low bulk DO conditions in the IFAS biofilm (**Figure 3C**). Compared with DO = 6 mg/L, expression under DO = 2 mg/L increased significantly for *acdA* (1.62-fold, *p* < 0.01), *acdB* (1.61-fold, *p* < 0.05), *ackA* (1.69-fold, *p* < 0.01), *pta* (1.86-fold, *p* < 0.001), *xfp* (1.61-fold, *p* < 0.01), and *por* (1.34-fold, *p* < 0.001). In contrast, only *acs* (1.15-fold) showed no significant change. These results indicate that DO conditions influenced not only nitrogen-cycling activity but also transcription of carbon-associated functions in *Ca.* Brocadia sapporoensis. In particular, the oxygen limitation conditions may preferentially enhance acetate transformation via *ackA*–*xfp* pathway and *ackA*–*pta* pathway in *Ca.* Brocadia sapporoensis. Together, these metagenomic and metatranscriptomic results support carbon metabolic flexibility in *Ca.* Brocadia sapporoensis and suggest that low-bulk-DO conditions in IFAS biofilms may preferentially enhance transcription of acetyl-phosphate-centered pathways.

### 3.3 Co-occurrence of High Total Nitrogen Loss and Ammonium Oxidation under Low Dissolved Oxygen Conditions

Motivated by genomic and transcriptomic evidence that low-DO conditions enhanced nitrogen- and carbon-associated functions in *Ca.* Brocadia sapporoensis within IFAS biofilm, two sets of aerobic batch assays were conducted across three bulk DO setpoints. Ammonia-fed assays were used to evaluate ammonium oxidation, NOx (nitrite plus nitrate) production, and TIN loss, whereas nitrite-fed assays were used as controls to evaluate nitrite oxidation and background NOx reduction under the same DO conditions. Together, these assays were designed to determine whether ammonium oxidation could be sustained alongside anammox-associated nitrogen loss under microaerobic conditions. In the ammonia-fed assays, the highest specific ammonia oxidation rate (sAOR) was achieved at a DO setpoint of 1.2 mg/L (4.13 ± 0.94 mg-N/g TS-h; **Figure 4A**). A comparable sAOR was observed at DO = 0.7 mg/L (3.94 ± 0.66 mg-N/g TS-h; **Figure 4A**), indicating that near-maximal ammonia oxidation capacity was maintained under moderately low DO conditions. In contrast, decreasing the DO concentration from 0.7 to 0.2 mg/L resulted in a 65.7% reduction in sAOR to 1.35 ± 0.14 mg-N/g TS-h (**Figure 4A**). One-way ANOVA followed by Tukey honest significant differences (Keselman and Rogan, 1977) post hoc test indicated that sAOR at DO = 0.2 mg/L was significantly lower than those at DO = 0.7 mg/L (*Tukey p* = 0.008) and DO = 1.2 mg/L (*Tukey p* = 0.005), whereas no significant difference was observed between DO = 0.7 and 1.2 mg/L (*Tukey p* = 0.936). The sharp decrease in ammonia oxidation rates at the lowest DO setpoint is likely attributable to oxygen limitation within the biofilm due mass transfer limitations (Regmi et al., 2011). Recent studies have shown that *Ca.* Nitrospira nitrosa, a comammox species previously detected in this same IFAS system (Cotto et al., 2023, 2020), exhibits an exceptionally low oxygen affinity constant (0.006 ± 0.001 mg-O_2_/L) (Hou et al., 2025). The observed DO–sAOR relationship in this study is therefore consistent with ammonia oxidation by nitrifiers adapted to low-oxygen conditions, while still being constrained at very low bulk DO due to biofilm-scale oxygen diffusion limitations. Notably, the ammonia oxidation rate at DO = 0.7 mg/L exceeded that reported for a recently studied AOB-dominated reactor operated under fully aerobic conditions (2.8–3.4 mg O_2_/L), which exhibited an ammonia oxidation rate of 3.15 mg-N/g VSS-h (Meng et al., 2025). This observation suggests that the nitrifier community in the IFAS-attached biomass collected from the aerobic zone (R4) may differ from conventional AOB-dominated suspended-growth systems and may include low-DO-adapted ammonia oxidizers capable of maintaining ammonium oxidation under microaerobic conditions. In previous analyses of the same full-scale IFAS system, kinetic and genomic evidence indicated that comammox bacteria were likely important aerobic ammonia oxidizers in the attached-growth phase (Cotto et al., 2023, 2020; Vilardi et al., 2023). Low-DO operational conditions have also been shown to promote shifts in nitrifier community structure from canonical AOB and NOB toward comammox bacteria (Xiang et al., 2025; Zheng et al., 2022), enabling low-DO systems to maintain nitrification rates comparable to those observed under conventional high-DO operation (Jimenez et al., 2026). However, the kinetic data alone cannot distinguish the relative contributions of AOB and comammox to ammonium oxidation. Therefore, combined with the comparable relative abundances of the recovered AOB and comammox MAGs observed in the IFAS biofilm, the measured sAOR should be interpreted as the combined activity of ammonia-oxidizing guilds within the attached biomass.

**Figure 4.**
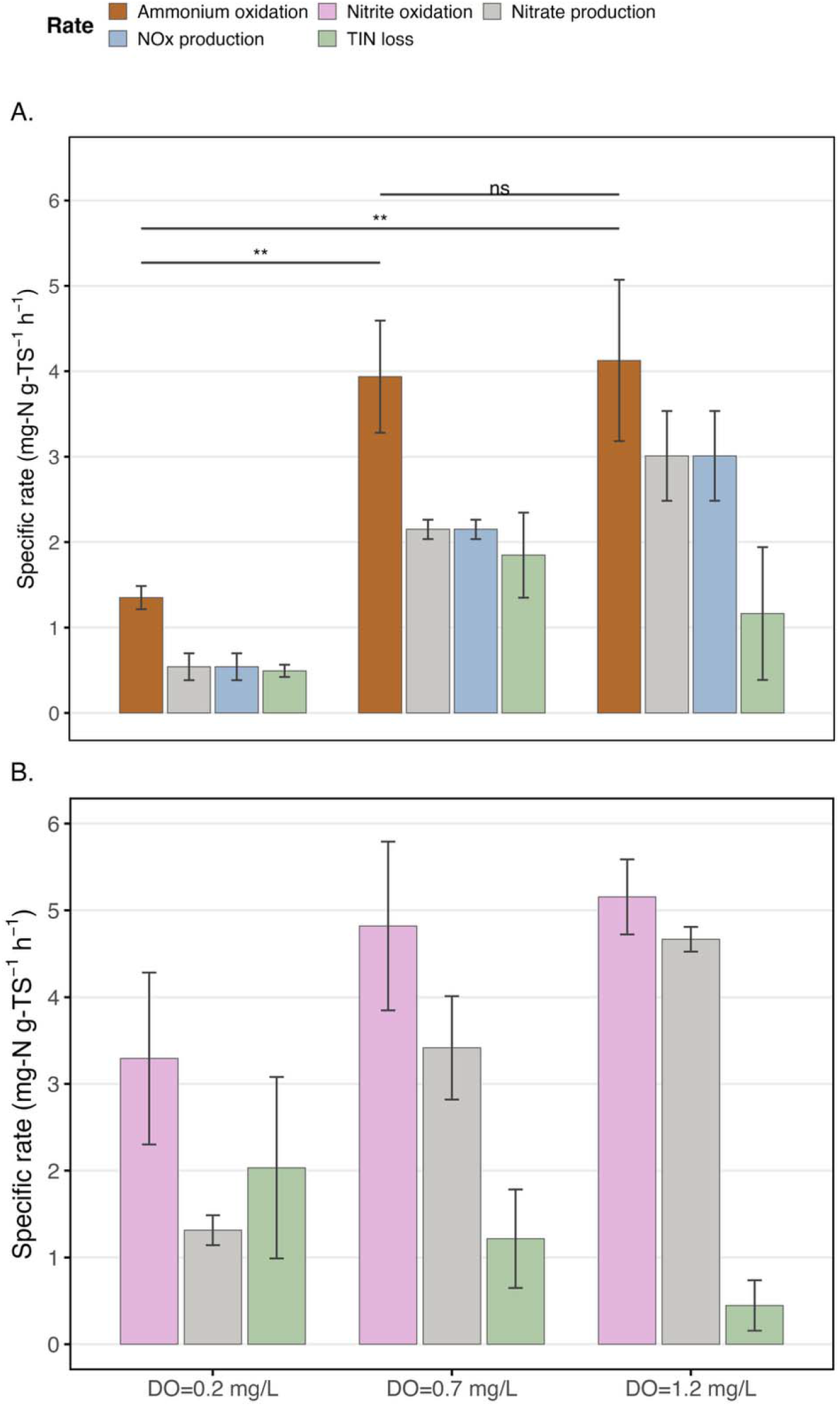
Aerobic nitrogen transformation rates in IFAS-attached biomass with an average attached TS concentration of 0.73 g TS/L. (A) Ammonia-fed assays initiated with NH_4_^+^-N for measuring ammonium oxidation, NOx production, nitrate production, and TIN loss. (B) Nitrite-fed control assays initiated with NO_2_^−^-N for measuring nitrite oxidation, nitrate production, and background TIN loss under the same DO setpoints.

In addition, no nitrite accumulation was observed throughout the batch tests operated at any of the three DO setpoints. Typically, maintaining low-oxygen conditions (0.1–1.5 mg/L) is considered a key operational strategy for achieving partial nitrification with nitrite accumulation (Y. Cao et al., 2017; Zuo et al., 2020), particularly in biofilm systems where competition among nitrifiers and oxygen concentration gradients within the biofilm favor nitrite buildup (Z. Wang et al., 2022; Winkler et al., 2011). However, under oxygen-limited conditions, nitrite produced via aerobic ammonia oxidation may be rapidly consumed via anammox, thereby preventing nitrite accumulation and directly contributing to nitrogen loss (Gottshall et al., 2021).

Measurable TIN loss was observed at all DO setpoints during aerobic ammonium oxidation assays, with the highest TIN loss rate occurring at DO = 0.7 mg/L (1.85 ± 0.50 mg-N/g TS-h), compared to lower rates at DO = 0.2 mg/L (0.49 ± 0.07 mg-N/g TS-h) and DO = 1.2 mg/L (1.16 ± 0.78 mg-N/g TS-h). To distinguish whether nitrogen loss in the ammonia-fed assays could be attributed solely to background NOx reduction under microaerobic conditions, nitrite-fed control assays were conducted at identical DO setpoints (**Figure 4B**). These control assays exhibited detectable TIN loss at all DO levels, confirming the presence of denitrification activity even under microaerobic conditions. VFA measurements confirmed the presence of background propionate (**Table S4**), indicating that a biodegradable fraction of the residual organic carbon in secondary effluent, which served as the batch test medium, may have supported this denitrification activity. However, under identical bulk DO conditions, aerobic ammonium oxidation assays consistently exhibited higher TIN loss rates than the corresponding nitrite oxidation controls at DO = 0.7 and 1.2 mg/L. Specifically, the additional TIN loss rate associated with the presence of ammonium was approximately 0.63 and 0.71 mg-N/g TS-h at DO = 0.7 and 1.2 mg/L, respectively, accounting for approximately 34% and 61% of the total TIN loss rates measured in the corresponding aerobic ammonium oxidation assays. Because the nitrite-oxidation controls quantify background NOx reduction under the same microaerobic conditions and medium, the additional nitrogen loss observed only when ammonium was present suggests an NH_4_^+^-linked N-removal pathway beyond background NOx reduction. Further support for this interpretation is provided by mass-balance closure between ammonium consumption, NOx production, and TIN loss. At DO = 0.7 and 1.2 mg/L, the deficit between ammonium consumption and NOx production closely matched the measured TIN loss, with relative deviations of only ∼3.2% and ∼4.1%, respectively. While microaerobic denitrification is clearly present, the excess TIN loss uniquely associated with ammonium is consistent with an additional N_2_-producing pathway that directly couples ammonium consumption to nitrogen loss. Together, these results are consistent with a contribution from anammox activity occurring within the IFAS biofilm under microaerobic conditions. Previous studies have shown that anammox enzymes can tolerate limited oxygen exposure and that irreversible inhibition generally occurs only at higher DO concentrations (>1.4 mg/L) (Egli et al., 2001; Yan et al., 2020). In contrast, no such enhancement in TIN loss was observed at DO = 0.2 mg/L, likely due to limited ammonium oxidation and insufficient nitrite availability to sustain detectable anammox activity.

### 3.4 Acetate amendment supports greater anammox-associated nitrogen loss than additional propionate amendment under anoxic conditions

The genomic and transcriptomic evidence indicated that the *Ca. Brocadia sapporoensis* MAG recovered from IFAS biofilm possesses the potential to reduce nitrate via a partial DNRA pathway, and that anammox activity contributed to a portion of the nitrogen loss observed under microaerobic conditions within the IFAS biofilm. Previous studies have also reported that certain anammox bacteria can utilize volatile fatty acids (VFAs), such as acetate and propionate, to support nitrate reduction via DNRA (Feng et al., 2019; Kartal et al., 2007b). Therefore, a series of anoxic batch tests was conducted in secondary effluent to evaluate how external acetate or propionate amendment affected nitrate reduction, nitrite availability, and downstream anammox-associated activity in this IFAS biofilm system. Because secondary effluent contained residual organic carbon, including background propionate averaging 17.92 mg COD/L (**Table S4**), these assays should be interpreted as testing the effect of additional VFA amendment on top of the wastewater-derived carbon matrix rather than comparing acetate and propionate as sole electron donors. The anammox activity in the absence of external VFAs was evaluated during batch test A (**Figure 5**). Anaerobic ammonium oxidation assays showed that the ratio of ammonium to nitrite consumption rates was approximately 1:1.06, which is close to the theoretical anammox stoichiometric ratio (1:1.146) (Lotti et al., 2014), supporting the occurrence of anammox activity. However, nitrate concentrations decreased during the batch test, with a measured nitrate consumption rate of 0.73 mg-N/g TS-h. This observation contradicts the canonical anammox stoichiometry, in which nitrate is produced as a byproduct (Lotti et al., 2014). In addition, the observed TIN loss rate (4.30 mg-N/g TS-h) substantially exceeded the theoretical nitrogen loss expected from anammox alone based on the measured ammonium consumption rate (1.64 mg-N/g TS-h). These results indicate that nitrogen removal under anaerobic conditions cannot be attributed solely to anammox. Instead, they suggest the coexistence of additional nitrate-reducing pathways, such as heterotrophic denitrification and/or DNRA, which likely consumed both the nitrate produced during anammox and the residual nitrate present at the beginning of the batch test. Previous studies have reported that certain anammox bacteria can use organic matter as an electron donor to convert nitrate into nitrite and ammonium, which could fuel the anammox process (Feng et al., 2019; Shu et al., 2016). Although no VFAs were externally added, the residual organic carbon in the secondary effluent could have served as an electron donor to support these nitrate reduction processes.

**Figure 5.**
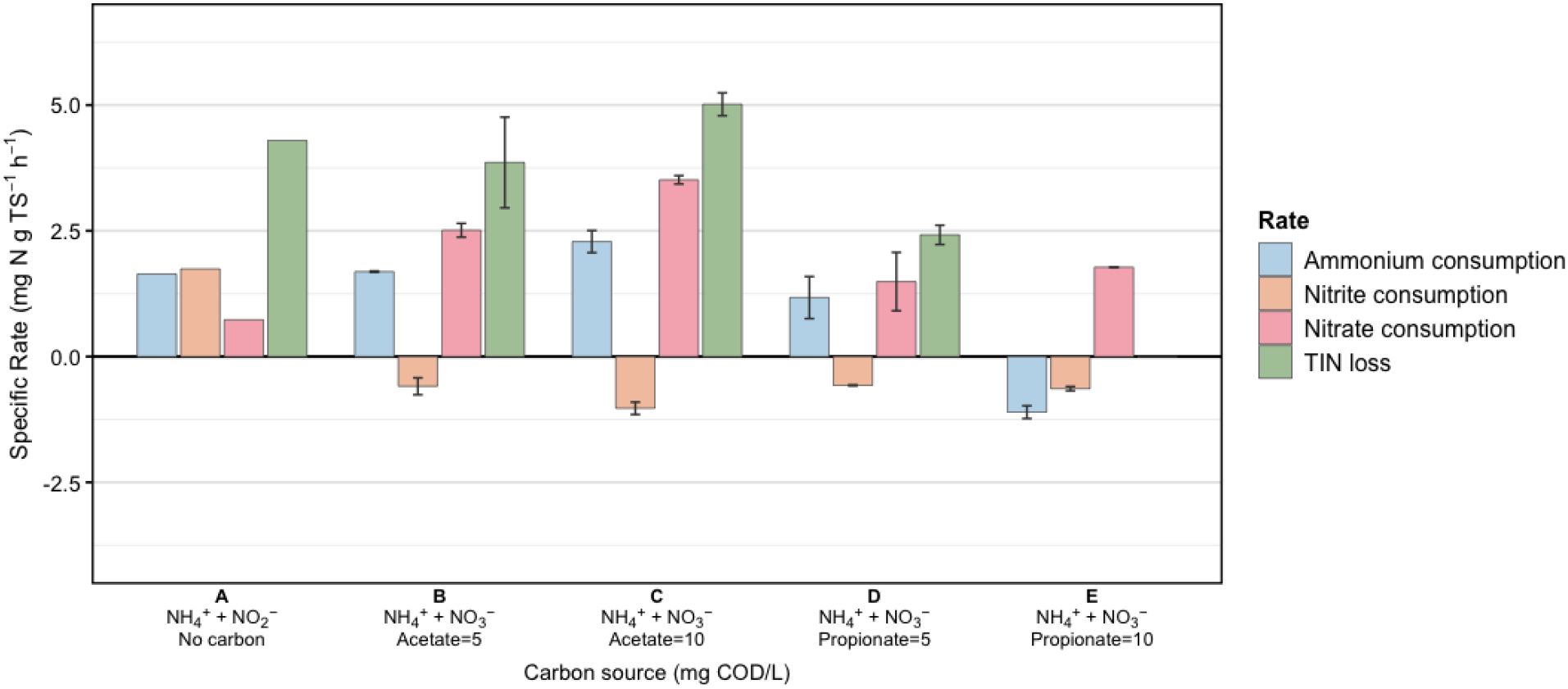
Effects of external acetate and propionate amendment on ammonium consumption, nitrite production, nitrate consumption, and TIN loss rates in anoxic batch assays using secondary effluent as the batch medium. The average attached TS concentration was 0.94 g TS/L.

Subsequently, batch tests B and C were conducted with external acetate amendment, nitrate, and ammonium under two acetate concentrations (5 and 10 mg COD/L, respectively) (**Figure 5**) to evaluate whether external acetate amendment could enhance nitrite availability through nitrate reduction and thereby support anammox-associated activity. The ammonium consumption rate was comparable between batch test B and batch test A but increased by 35.4% in batch test C where acetate concentrations were higher. Since nitrite was not externally supplied in batch tests B and C, the observed nitrite accumulation likely originated from nitrate reduction during the assay. Indeed, net nitrite accumulation was observed in both tests, corresponding to negative nitrite consumption (i.e., nitrite production) rates of –0.59 ± 0.17 and –1.03 ± 0.12 mg-N/g TS-h in batch tests B and C, respectively. Previous studies have demonstrated that low COD/NO_3_^−^–N ratios favor partial denitrification (S. Cao et al., 2017) and can promote partial DNRA–anammox coupling (Castro-Barros et al., 2017). In addition, the presence of ammonium has been shown to enhance nitrate reduction by anammox bacteria, as externally available ammonium circumvents the need for ammonium production via DNRA, allowing the accumulated nitrite to be directly utilized for energy-yielding anaerobic ammonium oxidation (Castro-Barros et al., 2017; M. K. H. Winkler et al., 2012). However, at low COD/NO_3_^−^–N ratios, electron donor limitation can constrain even the reduction of nitrate to nitrite (theoretical partial DNRA/denitrification COD/NO_3_^−^–N ≈ 1.14 based on stoichiometry). This likely explains the lower nitrite production rate observed in batch test B compared to batch test C, where a higher acetate concentration provided sufficient reducing equivalents to sustain more active nitrate reduction while still maintaining carbon-limited conditions favorable for partial DNRA–anammox coupling. The TIN loss rates were 3.86 ± 0.90 and 5.02 ± 0.23 mg-N/g TS-h in batch tests B and C, respectively. Because these assays were conducted under strictly anoxic conditions, the observed ammonium oxidation was predominantly attributed to anammox activity. Based on anammox stoichiometry (Lotti et al., 2014), approximately 3.33 and 4.54 mg-N/g TS-h of the TIN loss in batch tests B and C, respectively, can be attributed to anammox. This corresponds to 86.3% and 90.4% of the total TIN loss, suggesting that anammox-associated nitrogen removal accounted for most of the measured TIN loss under this stoichiometric framework. Collectively, these results support that nitrate reduction to nitrite occurred under acetate-amended conditions and likely supplied nitrite for downstream anammox, although the relative contributions of partial DNRA and partial denitrification could not be resolved from kinetic data alone.

Batch tests D and E were conducted by adding ammonium, nitrate, and 5 or 10 mg COD/L propionate to secondary effluent used as the batch medium to evaluate the effect of additional propionate amendment on anammox-associated activity (**Figure 5**). Compared to batch test A, the ammonium consumption rate decreased by 25.1% in batch test D with 5 mg COD/L as propionate. Furthermore, the ammonium consumption rate became negative (−1.11 ± 0.13 mg-N/g TS-h) in batch test E with 10 mg COD/L as propionate, consistent with net ammonium production through DNRA. Net nitrite accumulation was also observed in both batch tests D and E, which were conducted under the same COD/NO_3_^−^–N ratios as batch tests B and C, respectively. However, the TIN loss rate in batch test D was substantially lower than those observed in batch tests A, B, and C, and TIN loss in batch test E was negligible. For batch test D, the theoretical TIN loss estimated from the ammonium consumption rate based on anammox stoichiometry was approximately 2.31 mg-N/g TS-h, accounting for ∼95.6% of the measured TIN loss. This stoichiometric estimate suggests that anammox-associated activity accounted for most of the measured TIN loss under this condition, although contributions from other nitrogen transformation pathways cannot be fully excluded. In batch test E, TIN loss was negligible, as the measured TIN loss rate was slightly negative (−0.066 ± 0.056 mg-N/g TS-h), likely within the range of analytical uncertainty. This suggests that nitrate reduction in batch test E was largely decoupled from inorganic nitrogen removal, with nitrate potentially accumulating as nitrite or being further reduced to ammonium through DNRA. Previous studies have demonstrated that mixotrophic anammox bacteria can utilize both acetate and propionate as electron donors for nitrate reduction because of their corresponding carbon metabolic capacities (Chen et al., 2023; Feng et al., 2019). However, different anammox genera exhibit distinct preferences for organic substrates. For example, *Ca.* Brocadia fulgida showed the highest specific acetate oxidation rate, whereas *Ca.* Anammoxoglobus propionicus exhibited the highest specific propionate oxidation rate (Kartal et al., 2013). Although *Ca.* Brocadia sapporoensis has been reported to possess a dedicated propionate metabolism pathway that may support energetically favorable propionate degradation (Qiao et al., 2025; Tersteegen et al., 1997), the present results suggest that propionate was less favorable than acetate for supporting partial DNRA–anammox in this system. Previous studies have also shown that propionate can preferentially stimulate heterotrophic denitrifiers, which may intensify competition for propionate and redirect nitrate reduction toward nitrogen transformation pathways, rather than anammox-coupled nitrogen removal, compared to systems fed acetate (Chen et al., 2023; Qiao et al., 2025). We also observed substantial increases in soluble COD in the VFA-amended anoxic batch tests (**Figure S2**). Additional control experiments indicated that this COD increase was likely associated with elevated free ammonia concentrations caused by ammonium addition and pH increases during continuous N_2_ sparging (**Figure S3**), which promoted soluble organic matter release from the attached biomass rather than reflecting incomplete acetate consumption (**Text S2**). Because the observed nitrogen transformation rates remained consistent with anammox-associated nitrogen removal and the estimated free ammonia concentrations were below levels expected to substantially inhibit anammox activity (Aktan et al., 2012; Jaroszynski et al., 2012), this COD release likely affected the soluble COD profile but did not alter the interpretation of nitrogen removal dynamics.

Overall, under the secondary-effluent matrix used in these assays, external acetate amendment supported substantially greater anammox-associated nitrogen loss than additional propionate amendment. These results indicate that acetate was more favorable than propionate addition for promoting partial nitrate reduction coupled with anammox under the tested anoxic IFAS batch conditions.

### 3.5 Two-stage microaerobic–anoxic operation enhances anammox-associated nitrogen removal through co-occurring ammonium oxidation, anammox, and acetate-driven partial nitrate reduction

Based on the kinetic results from the aerobic and anoxic batch tests presented in the previous sections, we hypothesized that nitrogen removal by IFAS biofilm could be optimized in a single reactor by maintaining a target bulk DO of 0.7 mg/L during the microaerobic stage, followed by an anoxic stage with external acetate amendment. Under these conditions, nitrogen removal was hypothesized to occur through coordinated aerobic ammonium oxidation, nitrate reduction, and anammox activity across the two stages. To test this hypothesis, several batch tests with different control groups were conducted (**Table 1**). The sAOR achieved during the microaerobic stage at DO = 0.7 mg/L (3.61 ± 0.81 mg-N/g TS-h; **Figure 6A**) was comparable to that observed in the batch tests described in Section 3.3 at the same DO level (3.94 ± 0.66 mg-N/g TS-h; **Figure 4A**), indicating that similar aerobic ammonium oxidation activity was maintained under these conditions. Nitrite accumulation during the microaerobic stage remained minimal, while the NOx production rate (1.47 ± 0.30 mg-N/g TS-h; **Figure 6A**) was nearly identical to the nitrate production rate (1.45 ± 0.29 mg-N/g TS-h; **Figure 6A**), suggesting that full nitrification still dominated under this relatively low DO condition. The TIN loss rate during the microaerobic stage (1.77 ± 0.58 mg-N/g TS-h; **Figure 6A**) was also comparable to that observed previously at DO = 0.7 mg/L (1.85 ± 0.50 mg-N/g TS-h; **Figure 4A**), suggesting that similar nitrogen removal activity was maintained. The anammox control group further supported the coexistence of anammox activity under these microaerobic conditions, as the observed ammonium-to-nitrite consumption ratio (1:1.17) (**Figure 6A**) was close to the theoretical anammox stoichiometric ratio reported in previous studies (Lotti et al., 2014). Together, these findings support the hypothesis that aerobic ammonium oxidation and anammox can co-occur within the IFAS biofilm under microaerobic conditions (e.g., DO = 0.7 mg/L), thereby contributing to TIN removal during the microaerobic stage.

**Figure 6.**
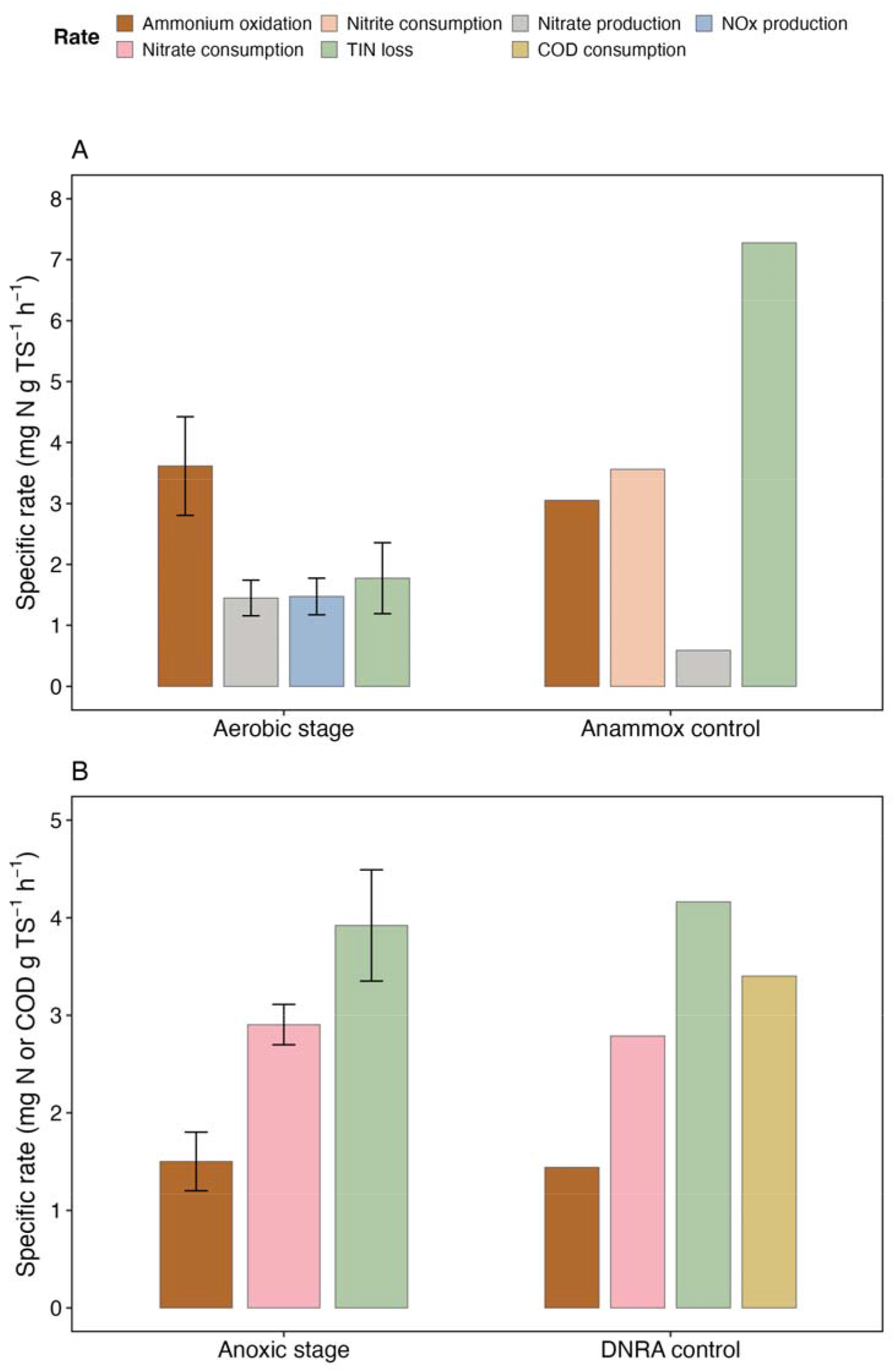
Nitrogen conversion kinetics during the two-stage microaerobic–anoxic batch tests. (A) Specific rates of nitrogen conversion during the microaerobic stage and anammox control group. (B) Specific rates of nitrogen and COD conversion during the anoxic stage and DNRA control group. The average attached TS concentration was 0.73 g TS/L.

During the anoxic stage, the sAOR was measured at 1.50 ± 0.30 mg-N/g TS-h (**Figure 6B**), suggesting that anammox remained active under these anoxic conditions. Nitrite accumulation was consistently observed during the first 60 min of the anoxic stage in all triplicate batch tests (**Figure S4A**), followed by a decline over the subsequent 60 min. The initial C/N ratio during the anoxic stage in all triplicate batch tests was 1.38 ± 0.04, which was comparable to the C/N ratios previously reported for partial denitrification/anammox systems (Gao et al., 2013; Sun et al., 2023). Because no significant nitrite accumulation was observed during the preceding microaerobic stage and no additional nitrite was added at the beginning of the anoxic stage, the nitrite supporting anammox activity was likely generated through nitrate reduction. Consistent with this interpretation, the nitrate consumption rate during the anoxic stage was 2.90 ± 0.21 mg-N/g TS-h, while the TIN removal rate reached 3.92 ± 0.57 mg-N/g TS-h (**Figure 6B**). Based on the measured sAOR and the stoichiometry of the anammox process, approximately 78% of the observed TIN removal was estimated to be attributable to anammox activity. Assuming that all nitrite consumed by anammox was supplied through nitrate reduction, the theoretical TIN removal rate calculated from anammox stoichiometry coupled with partial nitrate reduction was 3.99 mg-N/g TS-h. This value closely matched the measured TIN removal rate, further supporting that nitrate was partially reduced to nitrite during the anoxic stage, likely via partial DNRA and/or partial denitrification. In addition, approximately 9 mg COD/L was introduced through external acetate amendment at the beginning of the anoxic stage, while the total soluble COD concentration was approximately 49 mg/L because of residual organic carbon in the secondary effluent, including background VFAs such as propionate. COD was rapidly consumed during the first 40 min of the anoxic stage (**Figure S4B**), indicating that acetate and residual organic carbon likely served as electron donors for nitrate reduction. However, when the pH increased from an average of 7.5 to 7.7, COD increased again after 40 min of the anoxic stage, likely due to the same pH-associated mechanism discussed in Section 3.4.

In the DNRA control group amended with 10 mg COD/L acetate, 9 mg N/L NH_4_^+^, and 9 mg N/L NO_3_^−^, a comparable sAOR of 1.44 mg-N/g TS-h was observed (**Figure 6B**), suggesting that a similar level of anammox activity was maintained in this control relative to the anoxic stage. Nitrite accumulation was also observed during the first 60 min of the reaction period (**Figure S4A**), followed by gradual nitrite consumption during the remainder of the incubation. Based on the measurements, the ratio of COD consumption to nitrate reduction (rCOD/rNO_3_^−^-N) was 1.22, which was slightly higher than the stoichiometric rCOD/rNO_3_^−^-N ratio for partial DNRA/denitrification (1.14) but substantially lower than that required for full denitrification (2.86) reported in previous studies (Castro-Barros et al., 2017). These results further support that partial DNRA and/or partial denitrification occurred under the anoxic conditions of the DNRA control group, thereby supplying nitrite to sustain anammox-driven ammonium oxidation. In addition, the TIN removal rate in this control group was 4.16 mg-N/g TS-h, of which approximately 70.5% was estimated to be attributable to anammox activity based on the measured sAOR and the stoichiometry of the anammox process. Overall, these results support the hypothesis that nitrogen removal in the single-reactor system can be optimized by combining a microaerobic stage (DO = 0.7 mg/L) with a subsequent acetate-addition anoxic stage. Under these conditions, aerobic ammonium oxidation, nitrate reduction to nitrite, and anammox-associated activity likely acted in a coordinated manner across the two stages to enhance overall TIN removal.

### 3.6 Stage-specific transcriptional responses reveal complete nitrification and nitrate-reduction-supported anammox across microaerobic and acetate-amended anoxic stages

Endpoint TPM values were used to compare the transcriptional abundance of functional modules at the end of each stage, while DESeq2-derived log2 fold changes were used to determine the direction and magnitude of transcriptional responses, and Benjamini–Hochberg adjusted *p*-values were used to determine statistical significance. Complete DESeq2 results and endpoint TPM values for nitrogen- and carbon-metabolism genes are provided in **Tables S6**–S**8**. Based on the results, no significant changes in *amoABC* or *hao* expression were observed in either AOB or comammox bacteria during the microaerobic stage (**Figure 7**). The absence of upregulation may partly reflect the approximately 20-min pre-aeration period used to establish the target DO of 0.7 mg/L, during which transcription of the ammonia-oxidation genes may have already been upregulated. In addition, the sustained expression of *amoABC* and *hao* in both AOB and comammox bacteria based on the endpoint TPM values during the microaerobic stage (**Table S6**), together with the substantial aerobic ammonium oxidation rate measured in the batch tests suggests that both AOB and comammox likely contributed to ammonium oxidation during the microaerobic stage. For nitrite oxidation, canonical NOB *nxrAB* showed a statistically significant but quantitatively small increase of 1.07-fold (*padj* < 0.001), whereas comammox *nxrAB* showed no significant change. In contrast, comammox *nxrAB* was significantly upregulated when DO was decreased from 6 mg/L to 2 mg/L in our previous full-scale study (Johnston et al., 2024). Though a much lower DO of 0.7 mg/L was applied here, comammox *amoABC* and *hao* remained actively expressed without a corresponding increase in *nxrAB*. The minimal nitrite accumulation and close agreement between NOx and nitrate production rates, together with these transcriptional patterns, indicate that complete nitrification remained the dominant net process during the microaerobic stage. During the anoxic stage, *amoABC*, *hao*, and *nxrAB* in AOB, comammox, and canonical NOB generally showed nonsignificant changes due to the cessation of oxygen supply.

**Figure 7.**
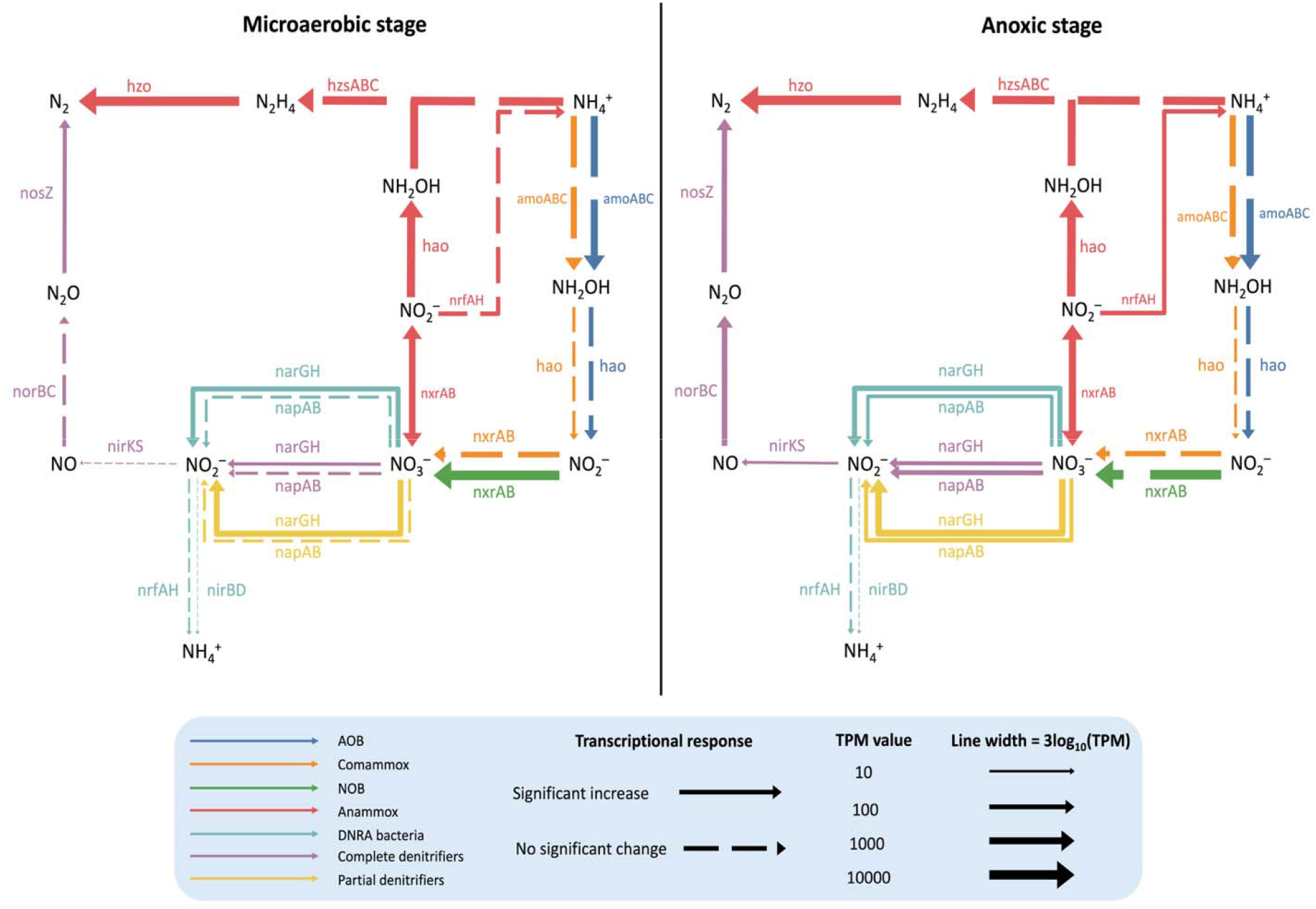
Nitrogen-transformation pathways and stage-specific transcriptional responses in IFAS biofilm during the two-stage batch test. Arrow colors indicate MAG groups, and arrow width represents endpoint expression levels scaled by log10-transformed mean TPM values. Gene labels indicate functional modules associated with each transformation step. Arrow line types indicate DESeq2-derived transcriptional responses from the beginning to the end of each stage: solid arrows indicate significant increases, and dashed arrows indicate no significant change. No significant decrease was observed for the pathways shown during either stage.

The expression of anammox-associated functional genes of *Ca.* Brocadia species (Lawson et al., 2017; Oshiki et al., 2016), including *hao*, *hzsABC*, and *hzo*, were also evaluated. The *hao* and *hzo* genes were significantly up-regulated by 1.47-fold (*padj* < 0.001) and 1.46-fold (*padj* < 0.001), respectively, during the microaerobic stage (**Figure 7; Table S6**). Their expression was also significantly elevated during the subsequent acetate-amended anoxic stage, increasing by 1.54-fold (*padj* < 0.001) and 1.24-fold (*padj* < 0.05), respectively. Nevertheless, the mean expression of *hzsABC* showed no significant change during both stages. Previous studies have shown that the *hzsABC* gene cluster is among the most highly expressed functional gene clusters in anammox bacteria (Kartal et al., 2011; Van De Vossenberg et al., 2013). In the present study, the average TPM of *hzsABC* at the beginning of the microaerobic stage was 5670.96, indicating that this gene cluster was already highly expressed before the batch experiment. Its persistently high TPM values at the aerobic and anoxic endpoints (**Table S6**) further suggest that the transcriptional potential for hydrazine synthesis from NH_4_^+^ and NH_2_OH was maintained during both stages (Oshiki et al., 2016), despite the absence of further transcriptional induction. Together, these transcriptional patterns are consistent with the batch-test results showing that anammox activity was maintained during the microaerobic stage and contributed to the observed TIN loss. This interpretation agrees with previous studies indicating that freshwater anammox bacteria, including *Ca.* Brocadia species, can tolerate low oxygen concentrations rather than behaving as obligate anaerobes under all conditions (Okabe et al., 2023; Zhang and Okabe, 2020). When considered together with the stoichiometric analysis, the sustained expression of core anammox-associated genes also supports the conclusion that anammox remained a major pathway contributing to nitrogen gas production during the subsequent anoxic stage. This interpretation is consistent with previous acetate-fed partial denitrification–anammox studies (Du et al., 2019; Wang et al., 2020). In addition, the relatively low VFA concentration supplied at the beginning of the anoxic stage did not appear to inhibit anammox activity, in agreement with previous studies (Shu et al., 2015).

More importantly, *nxrAB* in the *Ca.* Brocadia sapporoensis MAG was significantly upregulated by 2.28-fold (*padj* < 0.001) during the microaerobic stage (**Figure 7; Table S6**). In parallel, *narGH* was significantly upregulated in DNRA bacteria, complete denitrifiers, and partial denitrifiers by 2.20-fold (*padj* < 0.001), 2.43-fold (*padj* < 0.05), and 2.28-fold (*padj* < 0.001), respectively. In contrast, the downstream nitrite-reduction gene clusters, including *nrfAH* in the anammox and DNRA bacteria and *nirKS* in the complete denitrifiers, showed nonsignificant changes and low level of expression. These contrasting expression patterns together suggest that nitrate reduction to nitrite was more strongly stimulated than the subsequent reduction of nitrite to ammonium or nitric oxide during the microaerobic stage. Such preferential stimulation of nitrate reduction to nitrite may thereby providing an additional nitrite source for anammox during the microaerobic stage. Although *nxrAB* in anammox bacteria is generally proposed to catalyze nitrite oxidation to nitrate under aerobic conditions and contribute to nitrate overproduction (Sun et al., 2024), the catalyzed reaction is reversible (Kartal et al., 2007a). The significant upregulation of *nxrAB*, together with the substantially lower net nitrate production rate relative to the ammonium oxidation rate observed in the microaerobic-stage batch tests, therefore suggests that the *nxrAB* complex may also have operated in the nitrate-reducing direction under the tested microaerobic conditions. In addition, the secondary effluent used as the batch-test medium contained background propionate as described above. This residual propionate may have supplied carbon and reducing equivalents for nitrate reduction by anammox bacteria and heterotrophic nitrate reducers within oxygen-limited regions of the IFAS biofilm, although the anoxic amendment experiments indicated that additional propionate was less effective than external acetate amendment in enhancing anammox-associated TIN removal. The expression pattern of a single-stage partial nitritation–anammox–denitratation system operated under microaerobic and low-C/N conditions is consistent with the results observed in this study, where the combined expression of *napA* and *narG* was approximately threefold higher than that of *nirKS*, indicating preferential transcriptional investment in nitrate reduction over subsequent nitrite reduction (Zhou et al., 2022).

During the acetate-amended anoxic stage, *nxrAB* in *Ca*. Brocadia sapporoensis MAG was further significantly upregulated by 2.60-fold (*padj* < 0.001), whereas the downstream DNRA genes *nrfAH* showed a smaller but significant increase of 1.46-fold (*padj* < 0.05) (**Figure 7; Table S6**). Consistent with these differential responses, the endpoint TPM of *nxrAB* in *Ca*. Brocadia sapporoensis MAG was more than one order of magnitude higher than that of *nrfAH* (**Table S6**). A similar qualitative pattern was observed for the *Ca.* Brocadia sapporoensis MAG in the representative DNRA control, where both *nxrAB* and *nrfAH* expression increased, but endpoint expression of *nxrAB* remained substantially higher than that of *nrfAH* (**Figure S5C and Table S7**). Previous studies have shown that *nxr* can operate in the reverse direction to reduce nitrate to nitrite in anammox bacteria in the presence of organic and inorganic electron donors (Kartal et al., 2013). Under the anoxic, acetate-amended conditions applied here, reducing equivalents derived from acetate oxidation may therefore have favored operation of this *nxrAB* complex in the nitrate-reducing direction. Together with the kinetic and stoichiometric results described in Section 3.5, the stronger transcriptional response and substantially higher endpoint expression of *nxrAB* relative to *nrfAH* are consistent with the potential involvement of the anammox population in partial DNRA, in which nitrate was preferentially reduced to nitrite rather than fully reduced to ammonium and thus could be utilized by the anammox process (Castro-Barros et al., 2017; Kartal et al., 2007a). A similar nitrate-reduction response was observed in the other MAG groups. Both *narGH* and *napAB* were significantly upregulated in DNRA bacteria and in partial and complete denitrifiers, suggesting that these populations may also have contributed to nitrite production from nitrate. In addition, complete denitrifiers showed coordinated significant upregulation of *napAB*, *narGH*, *nirKS*, *norBC*, and *nosZ*, indicating activation of the complete denitrification pathway. Thus, complete denitrifiers may have competed with anammox for nitrite while also contributing to TIN removal and the reduction of denitrification intermediates, including N_2_O (Du et al., 2016; Cao et al., 2019). Nevertheless, the stoichiometric results, together with the substantially higher endpoint expression of *hao*, *hzsABC*, and *hzo* supports that anammox activity was the predominant pathway contributing to TIN removal during the acetate-amended anoxic stage. Again, the low C/N ratio of 1.38 ± 0.04 during the anoxic stage and 1.80 in the DNRA control may favor partial denitrification (Hou et al., 2023) and support anammox-associated DNRA (Castro-Barros et al., 2017), although the dominant response differs among anammox taxa (Chen et al., 2023; Feng et al., 2019; Shu et al., 2016). Thus, the system during the anoxic stage likely involved parallel nitrate-reduction routes, with partial DNRA and partial denitrification supplying nitrite to anammox while complete denitrification provided an additional, but smaller, nitrogen-removal pathway.

Carbon metabolism-related genes in the *Ca*. Brocadia sapporoensis MAG showed their strongest and most coordinated transcriptional response during the microaerobic stage. Specifically, *acs*, *acdA*, *pta*, *xfp*, and *porABC* were significantly upregulated, whereas *acdB* and *ackA* also showed positive but nonsignificant changes (**Figure S6**). Although no external carbon source was added during the microaerobic stage, the secondary effluent used as the batch medium contained background propionate, as described above. Previous studies have demonstrated that some anammox bacteria can oxidize propionate (Güven et al., 2005; Kartal et al., 2007b; Tao et al., 2019), and recent genome-resolved evidence combined with findings described in section 3.2 together indicates that *Ca.* Brocadia sapporoensis obtain the potential to exhibit mixotrophic metabolism under propionate-fed conditions (Qiao et al., 2025). Therefore, residual propionate may have contributed to the activation of propionyl-CoA metabolism, which in turn indirectly regulates other metabolic pathways, analogous to the role of acetate. In contrast, after 10 mg COD/L external acetate was added at the beginning of the anoxic stage, the carbon-metabolism genes showed only modest and variable changes. *acs*, *acdB*, *ackA*, and *xfp* exhibited positive mean changes, whereas *acdA*, *pta*, and *porABC* decreased (**Figure S6B; Table S8**), but none remained significant after multiple-testing correction, indicating that the short-term transcriptional responses were variable across biological replicates. Because the biomass had already been exposed to background propionate during the preceding microaerobic stage, the relevant carbon-metabolism pathways may already have been transcriptionally active before acetate addition, reducing the magnitude of the short-term incremental response. Despite the limited fold changes during the anoxic stage, xfp and ackA had the highest endpoint TPM values, followed by acs (**Figure S6A**; **Table S8**). This pattern indicates sustained transcriptional activity around the acetyl-phosphate node (Papini et al., 2012), while the expression of acs supports the simultaneous presence of a direct acetyl-CoA-forming acetate-activation route(Starai and Escalante-Semerena, 2004). In addition, *xfp* can connect pentose phosphate metabolism with acetyl-phosphate production through the cleavage of xylulose-5-phosphate (Shi et al., 2022), thereby potentially linking sugar-phosphate metabolism with acetyl-CoA-related central metabolism.

Overall, these expression results support the potential occurrence of anammox-associated partial DNRA–anammox coupling during both the microaerobic and acetate-amended anoxic stages. However, the present results cannot quantitatively resolve the relative contribution of anammox bacteria to nitrite production, because *narGH* and/or *napAB* were also significantly upregulated in DNRA bacteria, complete denitrifiers, and partial denitrifiers during both stages. In addition, complete denitrifiers showed coordinated upregulation of the full denitrification pathway during the anoxic stage, indicating that heterotrophic nitrate reducers may have competed with anammox bacteria for nitrate and accumulated nitrite while also contributing to nitrogen loss. Previous studies have reported that low C/N conditions favor both partial denitrification (Al-Hazmi et al., 2023) and anammox-associated DNRA (Castro-Barros et al., 2017), and that different carbon sources can have distinct effects on mixotrophic anammox process (Shu et al., 2015) and heterotrophic denitrification process (Zhang et al., 2024). Because all batch assays were conducted in secondary effluent containing residual organic carbon, the effects of acetate and propionate amendment should be interpreted within this wastewater-derived carbon background rather than as isolated single-carbon-source responses. Therefore, future long-term reactor studies should combine broader testing of wastewater-relevant VFAs across controlled concentration gradients under defined-carbon-source conditions and isotope-tracing techniques (Li et al., 2019; Trinh et al., 2025) to quantitatively distinguish nitrogen transformation pathways mediated by different groups of microorganisms within the same biofilm system and to identify the carbon source and loading range that best promotes anammox-coupled nitrogen removal.

## Conclusion

This study demonstrates that low-DO IFAS biofilms can support coupled nitrification, nitrate reduction, and anammox-based nitrogen removal. Metatranscriptomic analyses indicated that that low DO selectively enhanced nitrogen- and carbon-associated functions in the biofilm phase, and batch tests confirmed that ammonium oxidation and anammox could potentially co-occur under microaerobic conditions. Under anoxic conditions, VFA addition, particularly acetate, promoted partial nitrate reduction and enhanced anammox-associated nitrogen loss, indicating that nitrate-derived nitrite can support downstream anammox activity. Stage-specific transcriptional responses further suggested that *Ca.* Brocadia sapporoensis may contribute to its own nitrite supply through *nxrAB*-mediated partial nitrate reduction during both microaerobic and acetate-amended anoxic stages. Together, these findings highlight a potential operational strategy for mainstream nitrogen removal by coupling microaerobic ammonium oxidation with acetate-supported partial nitrate reduction and anammox in IFAS biofilms.

## Acknowledgements

This work was supported by the National Science Foundation (CBET-2428375) and by the Hampton Roads Sanitation District.

## Data availability

The raw sequencing data and anammox MAG are available in the NCBI Sequence Read Archive under BioProject accession PRJNA1504227.

## Reference

Aktan, C.K., Yapsakli, K., Mertoglu, B., 2012. Inhibitory effects of free ammonia on Anammox bacteria. Biodegradation 23, 751–762. 10.1007/s10532-012-9550-0

Al-Hazmi, H.E., Maktabifard, M., Grubba, D., Majtacz, J., Hassan, G.K., Lu, X., Piechota, G., Mannina, G., Bott, C.B., Mąkinia, J., 2023. An advanced synergy of partial denitrification-anammox for optimizing nitrogen removal from wastewater: A review. Bioresource Technology 381, 129168. 10.1016/j.biortech.2023.129168

Ali, T.U., 2013. Selective Inhibition of Ammonia Oxidation and Nitrite Oxidation Linked to N2O Emission with Activated Sludge and Enriched Nitrifiers. J. Microbiol. Biotechnol. 23, 719–723. 10.4014/jmb.1302.02017

American Public Health Association and others (Ed.), 2005. Standard methods for the examination of water & wastewater. APHA., 21st Ed. ed. American Public Health Association, Washington, DC.

Annavajhala, M.K., Kapoor, V., Santo-Domingo, J., Chandran, K., 2018. Comammox Functionality Identified in Diverse Engineered Biological Wastewater Treatment Systems. Environ. Sci. Technol. Lett. 5, 110–116. 10.1021/acs.estlett.7b00577

Bachmann, M., Lawrence, C., Wieczorek, N., Scott, T., Shelton, E., Elliott, B., Parsons, M., Klaus, S., Bott, C., 2025. Full-scale implementation of partial denitrification-anammox in IFAS processes: Cost savings and operational strategies. Water Environment Research 97, e70093. 10.1002/wer.70093

Bouras, G., Judd, L.M., Edwards, R.A., Vreugde, S., Stinear, T.P., Wick, R.R., 2024. How low can you go? Short-read polishing of Oxford Nanopore bacterial genome assemblies. Microbial Genomics 10. 10.1099/mgen.0.001254

Buakaew, T., Ratanatamskul, C., 2023. Effects of microaeration and sludge recirculation on VFA and nitrogen removal, membrane fouling reduction and microbial community of the anaerobic baffled biofilm-membrane bioreactor in treating building wastewater. Science of The Total Environment 903, 166248. 10.1016/j.scitotenv.2023.166248

Cao, S., Du, R., Li, B., Wang, S., Ren, N., Peng, Y., 2017. Nitrite production from partial-denitrification process fed with low carbon/nitrogen (C/N) domestic wastewater: performance, kinetics and microbial community. Chemical Engineering Journal 326, 1186–1196.

Cao, S., Du, R., Peng, Y., Li, B., Wang, S., 2019. Novel two stage partial denitrification (PD)-Anammox process for tertiary nitrogen removal from low carbon/nitrogen (C/N) municipal sewage. Chemical Engineering Journal 362, 107–115. 10.1016/j.cej.2018.12.160

Cao, Y., Van Loosdrecht, M.C.M., Daigger, G.T., 2017. Mainstream partial nitritation–anammox in municipal wastewater treatment: status, bottlenecks, and further studies. Appl Microbiol Biotechnol 101, 1365–1383. 10.1007/s00253-016-8058-7

Castro-Barros, C.M., Jia, M., Van Loosdrecht, M.C.M., Volcke, E.I.P., Winkler, M.K.H., 2017. Evaluating the potential for dissimilatory nitrate reduction by anammox bacteria for municipal wastewater treatment. Bioresource Technology 233, 363–372. 10.1016/j.biortech.2017.02.063

Chaumeil, P.-A., Mussig, A.J., Hugenholtz, P., Parks, D.H., 2020. GTDB-Tk: a toolkit to classify genomes with the Genome Taxonomy Database. Bioinformatics 36, 1925–1927. 10.1093/bioinformatics/btz848

Chen, C., Sun, F., Zhang, H., Wang, J., Shen, Y., Liang, X., 2016. Evaluation of COD effect on anammox process and microbial communities in the anaerobic baffled reactor (ABR). Bioresource Technology 216, 571–578. 10.1016/j.biortech.2016.05.115

Chen, H., Li, X., Liu, G., Zhu, J., Ma, X., Piao, C., You, S., Wang, K., 2023. Decoding the carbon and nitrogen metabolism mechanism in anammox system treating pharmaceutical wastewater with varying COD/N ratios through metagenomic analysis. Chemical Engineering Journal 457, 141316. 10.1016/j.cej.2023.141316

Chen, S., 2023. Ultrafast one-pass FASTQ data preprocessing, quality control, and deduplication using fastp. iMeta 2, e107. 10.1002/imt2.107

Chklovski, A., Parks, D.H., Woodcroft, B.J., Tyson, G.W., 2023. CheckM2: a rapid, scalable and accurate tool for assessing microbial genome quality using machine learning. Nat Methods 20, 1203–1212. 10.1038/s41592-023-01940-w

Cotto, I., Dai, Z., Huo, L., Anderson, C.L., Vilardi, K.J., Ijaz, U., Khunjar, W., Wilson, C., De Clippeleir, H., Gilmore, K., Bailey, E., Pinto, A.J., 2020. Long solids retention times and attached growth phase favor prevalence of comammox bacteria in nitrogen removal systems. Water Research 169, 115268. 10.1016/j.watres.2019.115268

Cotto, I., Vilardi, K.J., Huo, L., Fogarty, E.C., Khunjar, W., Wilson, C., De Clippeleir, H., Gilmore, K., Bailey, E., Lücker, S., Pinto, A.J., 2023. Low diversity and microdiversity of comammox bacteria in wastewater systems suggest specific adaptations within the Ca. Nitrospira nitrosa cluster. Water Research 229, 119497. 10.1016/j.watres.2022.119497

Cui, H., Zhang, L., Zhang, Q., Li, X., Peng, Y., 2023. Enrichment of comammox bacteria in anammox-dominated low-strength wastewater treatment system within microaerobic conditions: Cooperative effect driving enhanced nitrogen removal. Chemical Engineering Journal 453, 139851. 10.1016/j.cej.2022.139851

Daims, H., Lebedeva, E.V., Pjevac, P., Han, P., Herbold, C., Albertsen, M., Jehmlich, N., Palatinszky, M., Vierheilig, J., Bulaev, A., Kirkegaard, R.H., Von Bergen, M., Rattei, T., Bendinger, B., Nielsen, P.H., Wagner, M., 2015. Complete nitrification by Nitrospira bacteria. Nature 528, 504–509. 10.1038/nature16461

Danecek, P., Bonfield, J.K., Liddle, J., Marshall, J., Ohan, V., Pollard, M.O., Whitwham, A., Keane, T., McCarthy, S.A., Davies, R.M., Li, H., 2021. Twelve years of SAMtools and BCFtools. GigaScience 10, giab008. 10.1093/gigascience/giab008

De Coster, W., Rademakers, R., 2023. NanoPack2: population-scale evaluation of long-read sequencing data. Bioinformatics 39, btad311. 10.1093/bioinformatics/btad311

Dittrich, C.R., Bennett, G.N., San, K.-Y., 2008. Characterization of the Acetate-Producing Pathways in Escherichia coli. Biotechnol Progress 21, 1062–1067. 10.1021/bp050073s

Du, R., Cao, S., Li, B., Zhang, H., Wang, S., Peng, Y., 2019. Synergy of partial-denitrification and anammox in continuously fed upflow sludge blanket reactor for simultaneous nitrate and ammonia removal at room temperature. Bioresource Technology 274, 386–394. 10.1016/j.biortech.2018.11.101

Du, R., Cao, S., Wang, S., Niu, M., Peng, Y., 2016. Performance of partial denitrification (PD)-ANAMMOX process in simultaneously treating nitrate and low C/N domestic wastewater at low temperature. Bioresource Technology 219, 420–429. 10.1016/j.biortech.2016.07.101

Egli, K., Fanger, U., Alvarez, P.J.J., Siegrist, H., Van Der Meer, J.R., Zehnder, A.J.B., 2001. Enrichment and characterization of an anammox bacterium from a rotating biological contactor treating ammonium-rich leachate. Archives of Microbiology 175, 198–207. 10.1007/s002030100255

Feng, Y., Zhao, Y., Jiang, B., Zhao, H., Wang, Q., Liu, S., 2019. Discrepant gene functional potential and cross-feedings of anammox bacteria Ca. Jettenia caeni and Ca. Brocadia sinica in response to acetate. Water Research 165, 114974. 10.1016/j.watres.2019.114974

Gao, F., Zhang, H., Yang, F., Qiang, H., Li, H., Zhang, R., 2013. Study of an innovative anaerobic (A)/oxic (O)/anaerobic (A) bioreactor based on denitrification–anammox technology treating low C/N municipal sewage. Chemical Engineering Journal 232, 65–73. 10.1016/j.cej.2013.07.070

Ginestet, P., Audic, J.-M., Urbain, V., Block, J.-C., 1998. Estimation of Nitrifying Bacterial Activities by Measuring Oxygen Uptake in the Presence of the Metabolic Inhibitors Allylthiourea and Azide. Appl Environ Microbiol 64, 2266–2268. 10.1128/AEM.64.6.2266-2268.1998

Gottshall, E.Y., Bryson, S.J., Cogert, K.I., Landreau, M., Sedlacek, C.J., Stahl, D.A., Daims, H., Winkler, M., 2021. Sustained nitrogen loss in a symbiotic association of Comammox Nitrospira and Anammox bacteria. Water Research 202, 117426. 10.1016/j.watres.2021.117426

Güven, D., Dapena, A., Kartal, B., Schmid, M.C., Maas, B., Van De Pas-Schoonen, K., Sozen, S., Mendez, R., Op Den Camp, H.J.M., Jetten, M.S.M., Strous, M., Schmidt, I., 2005. Propionate Oxidation by and Methanol Inhibition of Anaerobic Ammonium-Oxidizing Bacteria. Appl Environ Microbiol 71, 1066–1071. 10.1128/AEM.71.2.1066-1071.2005

Hou, J., Zhu, Y., Shi, Y., Lin, L., Meng, F., Xu, M., Yang, L., Ni, B.-J., Chen, X., 2025. Enrichment and Kinetic Profiling of *Candidatus* Nitrospira nitrosa Culture Reveal Mechanisms Underlying Its Prevalence in Wastewater Treatment Systems. Environ. Sci. Technol. 59, 15272–15281. 10.1021/acs.est.5c04519

Hou, Z., Dong, W., Wang, H., Zhao, Z., Li, Z., Liu, H., Li, Y., Zeng, Z., Xie, J., Zhang, L., Liu, J., 2023. Response of nitrite accumulation to elevated C/NO– 3-N ratio during partial denitrification process: Insights of extracellular polymeric substance, microbial community and metabolic function. Bioresource Technology 384, 129269. 10.1016/j.biortech.2023.129269

Huang, X.-L., Gao, D.-W., Tao, Y., Wang, X.-L., 2014. C2/C3 fatty acid stress on anammox consortia dominated by Candidatus Jettenia asiatica. Chemical Engineering Journal 253, 402–407. 10.1016/j.cej.2014.05.055

Jaroszynski, L.W., Cicek, N., Sparling, R., Oleszkiewicz, J.A., 2012. Impact of free ammonia on anammox rates (anoxic ammonium oxidation) in a moving bed biofilm reactor. Chemosphere 88, 188–195. 10.1016/j.chemosphere.2012.02.085

Ji, X.-M., Zheng, C., Wang, Y.-L., Jin, R.-C., 2021. Decoding the interspecies interaction in anammox process with inorganic feeding through metagenomic and metatranscriptomic analysis. Journal of Cleaner Production 288, 125691. 10.1016/j.jclepro.2020.125691

Jimenez, J., Bauhs, K., Miller, M., Dold, P., Al-Omari, A., Garrido, M., Hiripitiyage, D., Wittman, M., Burger, G., Chandran, K., Sturm, B., 2026. Low dissolved oxygen nitrification through kinetic selection. Water Research 288, 124642. 10.1016/j.watres.2025.124642

Johnston, J., Vilardi, K., Cotto, I., Sudarshan, A., Bian, K., Klaus, S., Bachmann, M., Parsons, M., Wilson, C., Bott, C., Pinto, A., 2024. Metatranscriptomic Analysis Reveals Synergistic Activities of Comammox and Anammox Bacteria in Full-Scale Attached Growth Nitrogen Removal System. Environ. Sci. Technol. 58, 13023–13034. 10.1021/acs.est.4c04375

Kang, D.D., Li, F., Kirton, E., Thomas, A., Egan, R., An, H., Wang, Z., 2019. MetaBAT 2: an adaptive binning algorithm for robust and efficient genome reconstruction from metagenome assemblies. PeerJ 7, e7359. 10.7717/peerj.7359

Kartal, B., De Almeida, N.M., Maalcke, W.J., Op Den Camp, H.J.M., Jetten, M.S.M., Keltjens, J.T., 2013. How to make a living from anaerobic ammonium oxidation. FEMS Microbiol Rev 37, 428–461. 10.1111/1574-6976.12014

Kartal, B., Kuenen, J.G., Van Loosdrecht, M.C.M., 2010. Sewage Treatment with Anammox. Science 328, 702–703. 10.1126/science.1185941

Kartal, B., Kuypers, M.M.M., Lavik, G., Schalk, J., Op Den Camp, H.J.M., Jetten, M.S.M., Strous, M., 2007a. Anammox bacteria disguised as denitrifiers: nitrate reduction to dinitrogen gas via nitrite and ammonium. Environmental Microbiology 9, 635–642. 10.1111/j.1462-2920.2006.01183.x

Kartal, B., Maalcke, W.J., De Almeida, N.M., Cirpus, I., Gloerich, J., Geerts, W., Op Den Camp, H.J.M., Harhangi, H.R., Janssen-Megens, E.M., Francoijs, K.-J., Stunnenberg, H.G., Keltjens, J.T., Jetten, M.S.M., Strous, M., 2011. Molecular mechanism of anaerobic ammonium oxidation. Nature 479, 127–130. 10.1038/nature10453

Kartal, B., Rattray, J., Van Niftrik, L.A., Van De Vossenberg, J., Schmid, M.C., Webb, R.I., Schouten, S., Fuerst, J.A., Damsté, J.S., Jetten, M.S.M., Strous, M., 2007b. Candidatus “Anammoxoglobus propionicus” a new propionate oxidizing species of anaerobic ammonium oxidizing bacteria. Systematic and Applied Microbiology 30, 39–49. 10.1016/j.syapm.2006.03.004

Keselman, H.J., Rogan, J.C., 1977. The Tukey multiple comparison test: 1953–1976. Psychological Bulletin 84, 1050–1056. 10.1037/0033-2909.84.5.1050

Kits, K.D., Jung, M.-Y., Vierheilig, J., Pjevac, P., Sedlacek, C.J., Liu, S., Herbold, C., Stein, L.Y., Richter, A., Wissel, H., Brüggemann, N., Wagner, M., Daims, H., 2019. Low yield and abiotic origin of N2O formed by the complete nitrifier Nitrospira inopinata. Nat Commun 10, 1836. 10.1038/s41467-019-09790-x

Kits, K.D., Sedlacek, C.J., Lebedeva, E.V., Han, P., Bulaev, A., Pjevac, P., Daebeler, A., Romano, S., Albertsen, M., Stein, L.Y., Daims, H., Wagner, M., 2017. Kinetic analysis of a complete nitrifier reveals an oligotrophic lifestyle. Nature 549, 269–272. 10.1038/nature23679

Kleijn, R.J., Van Winden, W.A., Van Gulik, W.M., Heijnen, J.J., 2005. Revisiting the^13^ C-label distribution of the non-oxidative branch of the pentose phosphate pathway based upon kinetic and genetic evidence. The FEBS Journal 272, 4970–4982. 10.1111/j.1742-4658.2005.04907.x

Koch, H., Van Kessel, M.A.H.J., Lücker, S., 2019. Complete nitrification: insights into the ecophysiology of comammox Nitrospira. Appl Microbiol Biotechnol 103, 177–189. 10.1007/s00253-018-9486-3

Kolmogorov, M., Bickhart, D.M., Behsaz, B., Gurevich, A., Rayko, M., Shin, S.B., Kuhn, K., Yuan, J., Polevikov, E., Smith, T.P.L., Pevzner, P.A., 2020. metaFlye: scalable long-read metagenome assembly using repeat graphs. Nat Methods 17, 1103–1110. 10.1038/s41592-020-00971-x

Kopylova, E., Noé, L., Touzet, H., 2012. SortMeRNA: fast and accurate filtering of ribosomal RNAs in metatranscriptomic data. Bioinformatics 28, 3211–3217. 10.1093/bioinformatics/bts611

Krivoruchko, A., Zhang, Y., Siewers, V., Chen, Y., Nielsen, J., 2015. Microbial acetyl-CoA metabolism and metabolic engineering. Metabolic Engineering 28, 28–42. 10.1016/j.ymben.2014.11.009

Lawson, C.E., Nuijten, G.H.L., De Graaf, R.M., Jacobson, T.B., Pabst, M., Stevenson, D.M., Jetten, M.S.M., Noguera, D.R., McMahon, K.D., Amador-Noguez, D., Lücker, S., 2021. Autotrophic and mixotrophic metabolism of an anammox bacterium revealed by in vivo 13C and 2H metabolic network mapping. The ISME Journal 15, 673–687. 10.1038/s41396-020-00805-w

Lawson, C.E., Wu, S., Bhattacharjee, A.S., Hamilton, J.J., McMahon, K.D., Goel, R., Noguera, D.R., 2017. Metabolic network analysis reveals microbial community interactions in anammox granules. Nat Commun 8, 15416. 10.1038/ncomms15416

Li, H., 2018. Minimap2: pairwise alignment for nucleotide sequences. Bioinformatics 34, 3094–3100. 10.1093/bioinformatics/bty191

Li, J., Peng, Y., Zhang, L., Liu, J., Wang, X., Gao, R., Pang, L., Zhou, Y., 2019. Quantify the contribution of anammox for enhanced nitrogen removal through metagenomic analysis and mass balance in an anoxic moving bed biofilm reactor. Water Research 160, 178–187. 10.1016/j.watres.2019.05.070

Liang, Y., Li, D., Zhang, X., Zeng, H., Yang, Y., Zhang, J., 2015. Nitrate removal by organotrophic anaerobic ammonium oxidizing bacteria with C2/C3 fatty acid in upflow anaerobic sludge blanket reactors. Bioresource Technology 193, 408–414. 10.1016/j.biortech.2015.06.133

Lotti, T., Kleerebezem, R., Lubello, C., Van Loosdrecht, M.C.M., 2014. Physiological and kinetic characterization of a suspended cell anammox culture. Water Research 60, 1–14. 10.1016/j.watres.2014.04.017

Love, M.I., Huber, W., Anders, S., 2014. Moderated estimation of fold change and dispersion for RNA-seq data with DESeq2. Genome Biol 15, 550. 10.1186/s13059-014-0550-8

Martinez-Rabert, E., Smith, C.J., Sloan, W.T., Gonzalez-Cabaleiro, R., 2023. Competitive and substrate limited environments drive metabolic heterogeneity for comammox *Nitrospira*. ISME Communications 3, 91. 10.1038/s43705-023-00288-8

Meile, L., Rohr, L.M., Geissmann, T.A., Herensperger, M., Teuber, M., 2001. Characterization of the d -Xylulose 5-Phosphate/ d -Fructose 6-Phosphate Phosphoketolase Gene (*xfp*) from *Bifidobacterium lactis*. J Bacteriol 183, 2929–2936. 10.1128/JB.183.9.2929-2936.2001

Meng, Z., Yan, Y., Li, G., Li, Y., Wu, K., Zhang, Z., Reid, M.C., Gu, A.Z., 2025. New strategy for integration of anaerobic side-stream reactor with mainstream B-stage nitritation for short-cut nitrogen removal with granulation. Water Environment Research 97, e70056. 10.1002/wer.70056

Minh, B.Q., Schmidt, H.A., Chernomor, O., Schrempf, D., Woodhams, M.D., Von Haeseler, A., Lanfear, R., 2020. IQ-TREE 2: New Models and Efficient Methods for Phylogenetic Inference in the Genomic Era. Molecular Biology and Evolution 37, 1530–1534. 10.1093/molbev/msaa015

Nissen, J.N., Johansen, J., Allesøe, R.L., Sønderby, C.K., Armenteros, J.J.A., Grønbech, C.H., Jensen, L.J., Nielsen, H.B., Petersen, T.N., Winther, O., Rasmussen, S., 2021. Improved metagenome binning and assembly using deep variational autoencoders. Nat Biotechnol 39, 555–560. 10.1038/s41587-020-00777-4

Okabe, S., Ye, S., Lan, X., Nukada, K., Zhang, H., Kobayashi, K., Oshiki, M., 2023. Oxygen tolerance and detoxification mechanisms of highly enriched planktonic anaerobic ammonium-oxidizing (anammox) bacteria. ISME Communications 3, 45. 10.1038/s43705-023-00251-7

Oshiki, M., Ali, M., Shinyako-Hata, K., Satoh, H., Okabe, S., 2016. Hydroxylamine-dependent anaerobic ammonium oxidation (anammox) by “ *Candidatus* Brocadia sinica.” Environmental Microbiology 18, 3133–3143. 10.1111/1462-2920.13355

Pan, H., Yuan, D., Liu, W., Pi, Y., Wang, S., Zhu, G., 2020. Biogeographical distribution of dissimilatory nitrate reduction to ammonium (DNRA) bacteria in wetland ecosystems around the world. J Soils Sediments 20, 3769–3778. 10.1007/s11368-020-02707-y

Pan, S., Zhu, C., Zhao, X.-M., Coelho, L.P., 2022. A deep siamese neural network improves metagenome-assembled genomes in microbiome datasets across different environments. Nat Commun 13, 2326. 10.1038/s41467-022-29843-y

Papini, M., Nookaew, I., Siewers, V., Nielsen, J., 2012. Physiological characterization of recombinant Saccharomyces cerevisiae expressing the Aspergillus nidulans phosphoketolase pathway: validation of activity through 13C-based metabolic flux analysis. Appl Microbiol Biotechnol 95, 1001–1010. 10.1007/s00253-012-3936-0

Parks, D.H., Chuvochina, M., Waite, D.W., Rinke, C., Skarshewski, A., Chaumeil, P.-A., Hugenholtz, P., 2018. A standardized bacterial taxonomy based on genome phylogeny substantially revises the tree of life. Nat Biotechnol 36, 996–1004. 10.1038/nbt.4229

Pinto, A.J., Marcus, D.N., Ijaz, U.Z., Bautista-de Lose Santos, Q.M., Dick, G.J., Raskin, L., 2015. Metagenomic Evidence for the Presence of Comammox *Nitrospira* -Like Bacteria in a Drinking Water System. mSphere 1, e00054–15. 10.1128/mSphere.00054-15

Qiao, X., Zhang, Liyu, Yuan, T., Wu, Y., Geng, Y., Li, Y., Li, B., Zhang, Lijuan, Zhuang, W.-Q., Yu, K., 2025. Mixotrophic anammox bacteria outcompete dissimilatory nitrate reduction and denitrifying bacteria in propionate-containing wastewater. Bioresource Technology 419, 132077. 10.1016/j.biortech.2025.132077

R Core Team, 2021. R: A language and environment for statistical computing. R foundation for statistical computing, Vienna, Austria.

Ragsdale, S.W., 2003. Pyruvate Ferredoxin Oxidoreductase and Its Radical Intermediate. Chem. Rev. 103, 2333–2346. 10.1021/cr020423e

Regmi, P., Thomas, W., Schafran, G., Bott, C., Rutherford, B., Waltrip, D., 2011. Nitrogen removal assessment through nitrification rates and media biofilm accumulation in an IFAS process demonstration study. Water Research 45, 6699–6708. 10.1016/j.watres.2011.10.009

Roots, P., Wang, Y., Rosenthal, A.F., Griffin, J.S., Sabba, F., Petrovich, M., Yang, F., Kozak, J.A., Zhang, H., Wells, G.F., 2019. Comammox Nitrospira are the dominant ammonia oxidizers in a mainstream low dissolved oxygen nitrification reactor. Water Research 157, 396–405. 10.1016/j.watres.2019.03.060

Sakoula, D., Koch, H., Frank, J., Jetten, M.S.M., Van Kessel, M.A.H.J., Lücker, S., 2021. Enrichment and physiological characterization of a novel comammox *Nitrospira* indicates ammonium inhibition of complete nitrification. The ISME Journal 15, 1010–1024. 10.1038/s41396-020-00827-4

Schouten, S., Strous, M., Kuypers, M.M.M., Rijpstra, W.I.C., Baas, M., Schubert, C.J., Jetten, M.S.M., Sinninghe Damsté, J.S., 2004. Stable Carbon Isotopic Fractionations Associated with Inorganic Carbon Fixation by Anaerobic Ammonium-Oxidizing Bacteria. Appl Environ Microbiol 70, 3785–3788. 10.1128/AEM.70.6.3785-3788.2004

Schwengers, O., Jelonek, L., Dieckmann, M.A., Beyvers, S., Blom, J., Goesmann, A., 2021. Bakta: rapid and standardized annotation of bacterial genomes via alignment-free sequence identification: Find out more about Bakta, the motivation, challenges and applications, here. Microbial Genomics 7. 10.1099/mgen.0.000685

Shao, Y.-H., Wu, J.-H., 2021. Comammox *Nitrospira* Species Dominate in an Efficient Partial Nitrification–Anammox Bioreactor for Treating Ammonium at Low Loadings. Environ. Sci. Technol. 55, 2087–2098. 10.1021/acs.est.0c05777

Shao, Y.-H., Wu, J.-H., Chen, H.-W., 2024. Comammox Nitrospira cooperate with anammox bacteria in a partial nitritation–anammox membrane bioreactor treating low-strength ammonium wastewater at high loadings. Water Research 257, 121698. 10.1016/j.watres.2024.121698

Shi, L.-L., Zheng, Y., Tan, B.-W., Li, Z.-J., 2022. Establishment of a carbon-efficient xylulose cleavage pathway in Escherichia coli to metabolize xylose. Biochemical Engineering Journal 179, 108331. 10.1016/j.bej.2021.108331

Shu, D., He, Y., Yue, H., Gao, J., Wang, Q., Yang, S., 2016. Enhanced long-term nitrogen removal by organotrophic anammox bacteria under different C/N ratio constraints: quantitative molecular mechanism and microbial community dynamics. RSC Adv. 6, 87593–87606. 10.1039/C6RA04114K

Shu, D., He, Y., Yue, H., Zhu, L., Wang, Q., 2015. Metagenomic insights into the effects of volatile fatty acids on microbial community structures and functional genes in organotrophic anammox process. Bioresource Technology 196, 621–633. 10.1016/j.biortech.2015.07.107

Sieber, C.M.K., Probst, A.J., Sharrar, A., Thomas, B.C., Hess, M., Tringe, S.G., Banfield, J.F., 2018. Recovery of genomes from metagenomes via a dereplication, aggregation and scoring strategy. Nat Microbiol 3, 836–843. 10.1038/s41564-018-0171-1

Starai, V.J., Escalante-Semerena, J.C., 2004. Acetyl-coenzyme A synthetase (AMP forming). CMLS, Cell. Mol. Life Sci. 61. 10.1007/s00018-004-3448-x

Strous, M., Heijnen, J.J., Kuenen, J.G., Jetten, M.S.M., 1998. The sequencing batch reactor as a powerful tool for the study of slowly growing anaerobic ammonium-oxidizing microorganisms. Applied Microbiology and Biotechnology 50, 589–596. 10.1007/s002530051340

Sun, J., Feng, Y., Zheng, R., Wu, X., Kong, L., Zhang, K., Liu, S., 2024. Potential Growth of Anammox Bacteria under Aerobic Conditions. Environ. Sci. Technol. 58, 18244–18254. 10.1021/acs.est.4c06413

Sun, Y., Cao, J., Xu, R., Zhang, T., Luo, J., Xue, Z., Chen, S., Wang, S., Zhou, H., 2023. Influence of C/N ratio and ammonia on nitrogen removal and N2O emissions from one-stage partial denitrification coupled with anammox. Chemosphere 341, 140035. 10.1016/j.chemosphere.2023.140035

Tao, Y., Huang, X., Gao, D., Wang, X., Chen, C., Liang, H., Van Loosdrecht, M.C.M., 2019. NanoSIMS reveals unusual enrichment of acetate and propionate by an anammox consortium dominated by Jettenia asiatica. Water Research 159, 223–232. 10.1016/j.watres.2019.05.006

Tersteegen, A., Linder, D., Thauer, R.K., Hedderich, R., 1997. Structures and Functions of Four Anabolic 2-Oxoacid Oxidoreductases in *Methanobacterium Thermoautotrophicum*. European Journal of Biochemistry 244, 862–868. 10.1111/j.1432-1033.1997.00862.x

The Genome Standards Consortium, Bowers, R.M., Kyrpides, N.C., Stepanauskas, R., Harmon-Smith, M., Doud, D., Reddy, T.B.K., Schulz, F., Jarett, J., Rivers, A.R., Eloe-Fadrosh, E.A., Tringe, S.G., Ivanova, N.N., Copeland, A., Clum, A., Becraft, E.D., Malmstrom, R.R., Birren, B., Podar, M., Bork, P., Weinstock, G.M., Garrity, G.M., Dodsworth, J.A., Yooseph, S., Sutton, G., Glöckner, F.O., Gilbert, J.A., Nelson, W.C., Hallam, S.J., Jungbluth, S.P., Ettema, T.J.G., Tighe, S., Konstantinidis, K.T., Liu, W.-T., Baker, B.J., Rattei, T., Eisen, J.A., Hedlund, B., McMahon, K.D., Fierer, N., Knight, R., Finn, R., Cochrane, G., Karsch-Mizrachi, I., Tyson, G.W., Rinke, C., Lapidus, A., Meyer, F., Yilmaz, P., Parks, D.H., Murat Eren, A., Schriml, L., Banfield, J.F., Hugenholtz, P., Woyke, T., 2017. Minimum information about a single amplified genome (MISAG) and a metagenome-assembled genome (MIMAG) of bacteria and archaea. Nat Biotechnol 35, 725–731. 10.1038/nbt.3893

Trinh, H.P., Lee, S.-H., Ahn, J.H., Kim, D.-W., Gong, G., Park, H.-D., 2025. Insights into the one-stage DNRA-anammox reactors: Influence of carbon sources and C/N ratios. Chemical Engineering Journal 166959.

Van De Vossenberg, J., Woebken, D., Maalcke, W.J., Wessels, H.J.C.T., Dutilh, B.E., Kartal, B., Janssen-Megens, E.M., Roeselers, G., Yan, J., Speth, D., Gloerich, J., Geerts, W., Van Der Biezen, E., Pluk, W., Francoijs, K., Russ, L., Lam, P., Malfatti, S.A., Tringe, S.G., Haaijer, S.C.M., Op Den Camp, H.J.M., Stunnenberg, H.G., Amann, R., Kuypers, M.M.M., Jetten, M.S.M., 2013. The metagenome of the marine anammox bacterium ‘ *Candidatus* Scalindua profunda’ illustrates the versatility of this globally important nitrogen cycle bacterium. Environmental Microbiology 15, 1275–1289. 10.1111/j.1462-2920.2012.02774.x

Van Kessel, M.A.H.J., Speth, D.R., Albertsen, M., Nielsen, P.H., Op Den Camp, H.J.M., Kartal, B., Jetten, M.S.M., Lücker, S., 2015. Complete nitrification by a single microorganism. Nature 528, 555–559. 10.1038/nature16459

Vilardi, K., Cotto, I., Bachmann, M., Parsons, M., Klaus, S., Wilson, C., Bott, C.B., Pieper, K.J., Pinto, A.J., 2023. Co-Occurrence and Cooperation between Comammox and Anammox Bacteria in a Full-Scale Attached Growth Municipal Wastewater Treatment Process. Environ. Sci. Technol. 57, 5013–5023. 10.1021/acs.est.2c09223

Wang, D., Zheng, Q., Huang, K., Springael, D., Zhang, X.-X., 2020. Metagenomic and metatranscriptomic insights into the complex nitrogen metabolic pathways in a single-stage bioreactor coupling partial denitrification with anammox. Chemical Engineering Journal 398, 125653. 10.1016/j.cej.2020.125653

Wang, W., Wang, T., Liu, Q., Wang, H., Xue, H., Zhang, Z., Wang, Y., 2022. Biochar-mediated DNRA pathway of anammox bacteria under varying COD/N ratios. Water Research 212, 118100. 10.1016/j.watres.2022.118100

Wang, Z., Zheng, M., Duan, H., Yuan, Z., Hu, S., 2022. A 20-Year Journey of Partial Nitritation and Anammox (PN/A): from Sidestream toward Mainstream. Environ. Sci. Technol. 56, 7522–7531. 10.1021/acs.est.1c06107

Wick, R.R., Holt, K.E., 2022. Polypolish: Short-read polishing of long-read bacterial genome assemblies. PLoS Comput Biol 18, e1009802. 10.1371/journal.pcbi.1009802

Winkler, M.-K., Kleerebezem, R., Van Loosdrecht, M., 2012. Integration of anammox into the aerobic granular sludge process for main stream wastewater treatment at ambient temperatures. Water research 46, 136–144.

Winkler, M.K.H., Kleerebezem, R., Kuenen, J.G., Yang, J., Van Loosdrecht, M.C.M., 2011. Segregation of Biomass in Cyclic Anaerobic/Aerobic Granular Sludge Allows the Enrichment of Anaerobic Ammonium Oxidizing Bacteria at Low Temperatures. Environ. Sci. Technol. 45, 7330–7337. 10.1021/es201388t

Winkler, M.K.H., Yang, J., Kleerebezem, R., Plaza, E., Trela, J., Hultman, B., Van Loosdrecht, M.C.M., 2012. Nitrate reduction by organotrophic Anammox bacteria in a nitritation/anammox granular sludge and a moving bed biofilm reactor. Bioresource Technology 114, 217–223. 10.1016/j.biortech.2012.03.070

Woodcroft, B.J., Singleton, C.M., Boyd, J.A., Evans, P.N., Emerson, J.B., Zayed, A.A.F., Hoelzle, R.D., Lamberton, T.O., McCalley, C.K., Hodgkins, S.B., Wilson, R.M., Purvine, S.O., Nicora, C.D., Li, C., Frolking, S., Chanton, J.P., Crill, P.M., Saleska, S.R., Rich, V.I., Tyson, G.W., 2018. Genome-centric view of carbon processing in thawing permafrost. Nature 560, 49–54. 10.1038/s41586-018-0338-1

Xiang, Y., Song, X., Yang, Y., Deng, S., Fu, L., Yang, C., Chen, M., Pu, J., Zhang, H., Chai, H., 2025. Comammox rather than AOB dominated the efficient autotrophic nitrification-denitrification process in an extremely oxygen-limited environment. Water Research 268, 122572. 10.1016/j.watres.2024.122572

Xie, J., Chen, Y., Cai, G., Cai, R., Hu, Z., Wang, H., 2023. Tree Visualization By One Table (tvBOT): a web application for visualizing, modifying and annotating phylogenetic trees. Nucleic Acids Research 51, W587–W592. 10.1093/nar/gkad359

Xu, S., Chai, W., Xiao, R., Wang, B., Lu, H., 2023. Synergy between Comammox and Anammox Bacteria in Wastewater Ammonia Removal. ACS EST Eng. 3, 1582–1591. 10.1021/acsestengg.3c00144

Yan, Y., Wang, W., Wu, M., Jetten, M.S.M., Guo, J., Ma, J., Wang, H., Dai, X., Wang, Y., 2020. Transcriptomics Uncovers the Response of Anammox Bacteria to Dissolved Oxygen Inhibition and the Subsequent Recovery Mechanism. Environ. Sci. Technol. 54, 14674–14685. 10.1021/acs.est.0c02842

Yang, W., He, S., Han, M., Wang, B., Niu, Q., Xu, Y., Chen, Y., Wang, H., 2018. Nitrogen removal performance and microbial community structure in the start-up and substrate inhibition stages of an anammox reactor. Journal of Bioscience and Bioengineering 126, 88–95. 10.1016/j.jbiosc.2018.02.004

Yin, X., Rahaman, M.H., Liu, W., Mąkinia, J., Zhai, J., 2021. Comparison of nitrogen and VFA removal pathways in autotrophic and organotrophic anammox reactors. Environmental Research 197, 111065. 10.1016/j.envres.2021.111065

Zhang, L., Okabe, S., 2020. Ecological niche differentiation among anammox bacteria. Water Research 171, 115468. 10.1016/j.watres.2020.115468

Zhang, M., Liu, J., Wang, D., Lu, M., Fan, Y., Ji, J., Wu, J., 2024. Combined effects of carbon source and C/N ratio on the partial denitrification performance: Nitrite accumulation, denitrification kinetic and microbial transition. Journal of Environmental Chemical Engineering 12, 113343. 10.1016/j.jece.2024.113343

Zhang, S., Zhang, Z., Xia, S., Ding, N., Liao, X., Yang, R., Chen, M., Chen, S., 2021. The potential contributions to organic carbon utilization in a stable acetate-fed Anammox process under low nitrogen-loading rates. Science of The Total Environment 784, 147150. 10.1016/j.scitotenv.2021.147150

Zhao, Y., Jiang, B., Tang, X., Liu, S., 2019. Metagenomic insights into functional traits variation and coupling effects on the anammox community during reactor start-up. Science of The Total Environment 687, 50–60. 10.1016/j.scitotenv.2019.05.491

Zheng, M., Mu, G., Zhang, A., Wang, J., Chang, F., Niu, J., Wang, X., Gao, T., Zhao, Z., 2022. Predominance of comammox bacteria among ammonia oxidizers under low dissolved oxygen condition. Chemosphere 308, 136436. 10.1016/j.chemosphere.2022.136436

Zheng, M., Wang, M., Zhao, Z., Zhou, N., He, S., Liu, S., Wang, J., Wang, X., 2019. Transcriptional activity and diversity of comammox bacteria as a previously overlooked ammonia oxidizing prokaryote in full-scale wastewater treatment plants. Science of The Total Environment 656, 717–722. 10.1016/j.scitotenv.2018.11.435

Zhou, F., Xiao, W.-Z., Zhou, K.-Y., Zhuang, J.-L., Zhang, X., Liu, Y.-D., Ni, B.-J., Shapleigh, J.P., Zhou, M., Luo, X.-Z., Li, W., 2022. Performance characteristics and community analysis of a single-stage partial nitritation, anammox and denitratation (SPANADA) integrated process for treating low C/N ratio wastewater. Chemical Engineering Journal 433, 134452. 10.1016/j.cej.2021.134452

Zimin, A.V., Salzberg, S.L., 2020. The genome polishing tool POLCA makes fast and accurate corrections in genome assemblies. PLoS Comput Biol 16, e1007981. 10.1371/journal.pcbi.1007981

Zuo, F., Sui, Q., Zheng, R., Ren, J., Wei, Y., 2020. In situ startup of a full-scale combined partial nitritation and anammox process treating swine digestate by regulation of nitrite and dissolved oxygen. Bioresource Technology 315, 123837. 10.1016/j.biortech.2020.123837

