## Supplementary Information for "Genome-resolved and kinetic evidence for low-DO comammox–anammox synergy and acetate-stimulated nitrate reduction in IFAS biofilms"

Texts: 3

Figures: 6

Tables: 8

Page: 26

**Text S1. Library preparation and sequencing details for short- and long-read metagenomic sequencing**

For short-read sequencing, DNA extracts were prepared using the NEBNext Ultra II FS DNA Library Prep Kit and sequenced on the Element Biosciences AVITI and Illumina NovaSeq X platforms using 2 × 150 bp paired-end sequencing by the Molecular Evolution Core at the Parker H. Petit Institute for Bioengineering and Bioscience, Georgia Institute of Technology. Sequencing reads generated from both platforms were combined for downstream analyses. For long-read sequencing, DNA extracts were prepared using the Ligation Sequencing gDNA Native Barcoding Kit 24 V14 (SQK-NBD114.24, Oxford Nanopore Technologies) according to the manufacturer’s protocol for the R10.4.1 flow cell. Libraries were sequenced on an Oxford Nanopore MinION Mk1C platform.

**Text S2. Evaluation of soluble COD increases during anoxic VFA-amended batch tests**

Substantial increases in soluble COD were observed in the main VFA-amended anoxic batch tests described in Section 3.4. To determine whether this increase reflected incomplete VFA consumption or release of soluble organic matter from the attached biomass, six supplementary diagnostic control batch experiments were conducted without pH control under different substrate-addition and N_2_-sparging conditions. These controls were designed to identify operational conditions associated with pH increase, elevated free ammonia (FA), and COD release, rather than to quantify nitrogen transformation kinetics. Their initial substrate concentrations and operational conditions are summarized in **Table S2**, and the corresponding pH, FA, and COD responses are summarized in **Table S3**. The results showed that substantial increases in soluble COD were only observed in control groups with initial ammonium addition and concurrent increases in pH during the batch tests (**Table S3**). In contrast, control groups with ammonium addition and without continuous sparging of dinitrogen gas exhibited relatively stable pH and much lower FA concentrations and did not show significant increases in soluble COD, where the FA concentrations in these groups were calculated using equation {1.214 × [NH_4_^+^–N] × 10^pH^}/{exp[6344/(273 + T($℃$))] + 10^pH^} (Anthonisen et al., 1976). The pH increase observed during the batch tests could be partially attributed to alkalinity generation associated with nitrogen transformation processes such as denitrification, anammox, and DNRA. In addition, continuous sparging of dinitrogen gas at a relatively high flow rate was applied to maintain anaerobic conditions during the batch tests, which likely enhanced CO_2_ stripping from the liquid phase and further contributed to the observed pH increase. As FA concentration is strongly dependent on pH and increases exponentially with increasing pH under a given ammonium concentration and temperature, the elevated pH conditions substantially increased FA levels in these batch systems. VFA concentrations at the beginning and end of these control groups were measured (**Table S4**), where the added acetate was completely consumed during the batch tests when nitrate was present as electron acceptor, while high concentrations of butyrate were formed in the groups containing ammonium, which therefore exhibited large increases in COD. Previous studies have reported that FA can disrupt the recalcitrant structure of the sludge, leading to the release of extracellular polymeric substances (EPS) and intracellular materials from the solid phase into the liquid phase (Hejnfelt and Angelidaki, 2009; Yang et al., 2018), and promote the release of soluble organic matter like short-chain fatty acids during fermentation, leading to significant increases in soluble COD over time (Zhang et al., 2018; Zhao et al., 2018). Therefore, the formation of butyrate in these control groups may be associated with anaerobic fermentation pathways in mixed microbial communities, where EPS and intracellular storage compounds can be converted into longer-chain fatty acids under reducing conditions. To further examine the relationship between FA concentration and soluble COD release, additional batch experiments were conducted under controlled conditions (**Table S5**), where FA concentrations were adjusted by varying pH while maintaining the same ammonium concentrations. The results showed that the increase in soluble COD was positively correlated with FA concentration and followed a logarithmic relationship (R^2^ = 0.92) (**Figure S3**). These results further suggest that increasing FA concentrations can substantially enhance the release of soluble organic matter into the liquid phase. Previous studies also demonstrated that higher FA concentrations have been associated with greater release of soluble substrates, and the increase of FA even in a small range (0.48–1.5 mg/L) has been reported to facilitate sludge disintegration (Zhao et al., 2018). Generally, FA can diffuse freely through cell membranes and disrupt intracellular proton and potassium gradients without energy consumption, potentially leading to cell inactivation and structural destabilization (Kayhanian, 1999; Liu et al., 2019). Applying this interpretation to the main VFA-amended anoxic batch tests described in Section 3.4, the soluble COD increases observed in batch tests B–E were likely attributable to elevated FA concentrations resulting from ammonium addition and pH increases. The average final FA concentrations reached 4.12, 4.22, 4.09, and 5.12 mg/L in batch tests B, C, D, and E, respectively. Although FA has been reported to inhibit anammox activity at elevated concentrations, approximately 48.6 mg/L is generally required to achieve around 50% inhibition (Aktan et al., 2012; Jaroszynski et al., 2012), which is much higher than the FA concentrations observed in this study. Therefore, the FA levels observed in this study were more likely to contribute to biomass disintegration and soluble organic matter release rather than causing significant inhibition of anammox activity.

**Text S3. Control batch tests for evaluating soluble COD increases during anoxic VFA-amended assays**

Additional control batch tests were conducted to identify potential causes of soluble COD increases observed during selected anoxic VFA-amended assays. IFAS carrier media were collected from R4, transported submerged in secondary clarifier effluent, and shipped overnight to Georgia Institute of Technology. Before the assays, IFAS carrier media were gently washed twice with secondary clarifier effluent to remove loosely attached biomass and then transferred into 1 L Pyrex media bottles. Each bottle contained 30 carrier media pieces suspended in 800 mL of secondary clarifier effluent. For control groups operated with continuous N_2_ purging, bottles were fitted with two-port delivery caps. One port was connected to an ultrapure N_2_ gas supply to maintain anoxic conditions throughout the 3 h incubation, while the second port was left open to the atmosphere to maintain pressure equilibrium. For the control group without continuous N_2_ purging, bottles were purged with N_2_ for 30 min before substrate addition and then immediately sealed after substrate addition to maintain anoxic conditions during incubation. Six control batch tests were conducted with different initial substrate conditions to assess the effects of substrate addition, continuous N_2_ purging, pH changes, and abiotic transformations on soluble COD dynamics (**Table S2**). Substrates were injected using a syringe. Aqueous samples were collected immediately after substrate addition and after 3 h of incubation, filtered through 0.22 μm syringe filters, and analyzed for nitrogen species and soluble COD. pH and temperature were measured before substrate addition and at the end of each assay using a Thermo Scientific Orion Star A329 portable meter.

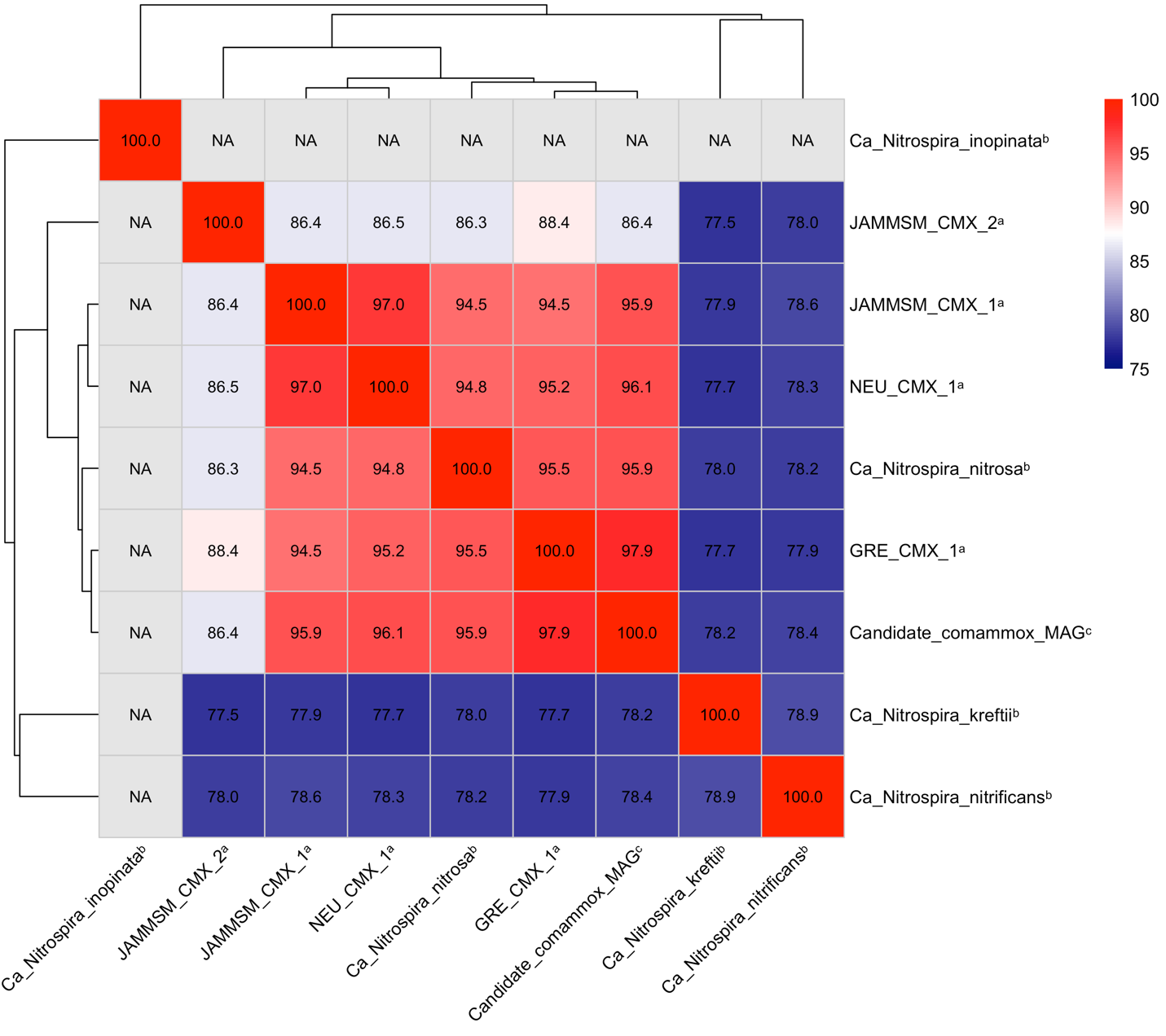

**Figure S1.** **FastANI-based average nucleotide identity heatmap comparing the candidate comammox MAG recovered in this study with previously recovered wastewater-derived comammox MAGs and NCBI reference genomes. Superscripts indicate genome sources: ^a^ Cotto et al. (2023); ^b^ NCBI reference genomes—*Ca.* N. kreftii (GCA_014058405.1), *Ca.* N. nitrificans (GCF_001458775.1), *Ca.* N. nitrosa (GCF_001458735.1), and *Ca.* N. inopinata (GCF_001458695.1); and ^c^ this study.**

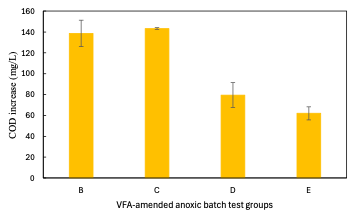

**Figure S2. Increase in soluble COD during VFA-amended anoxic batch tests.**

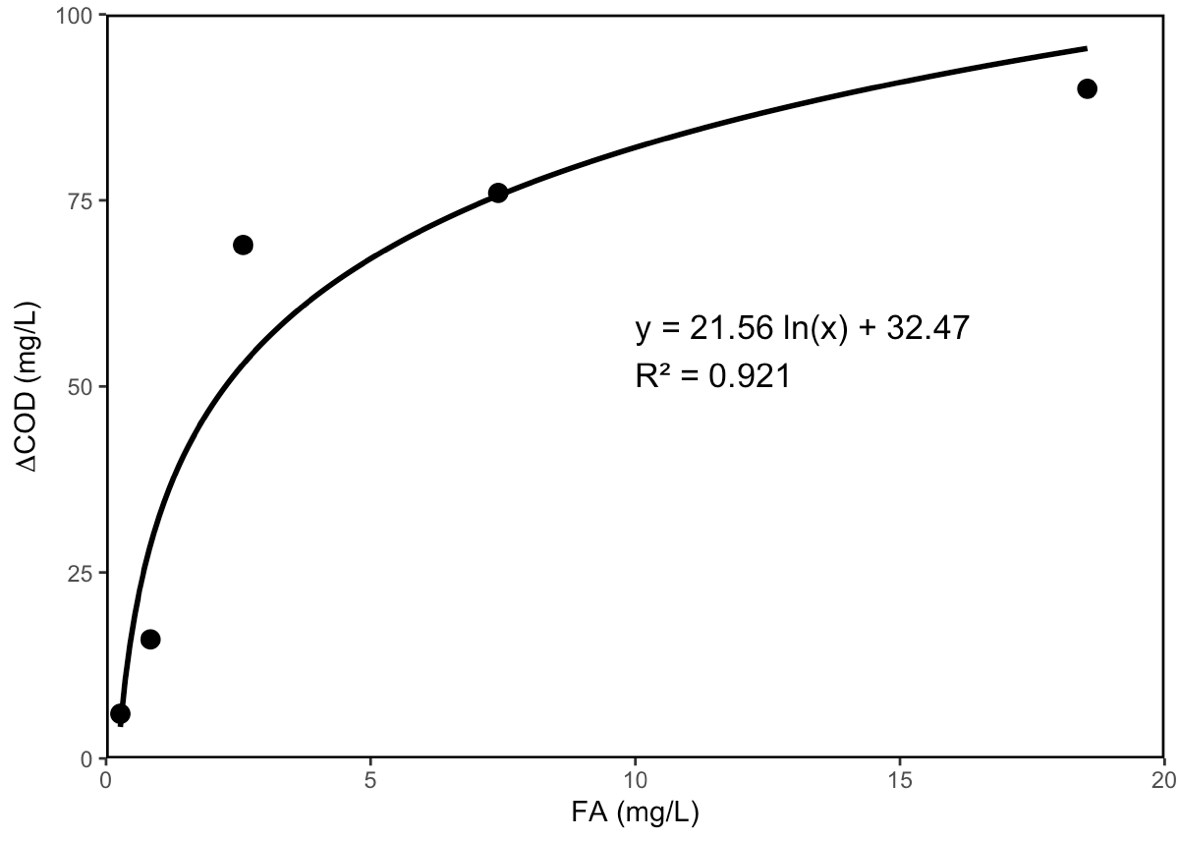

**Figure S3. Relationship between calculated free ammonia concentration and soluble COD release during supplementary adjusted-pH control tests.**

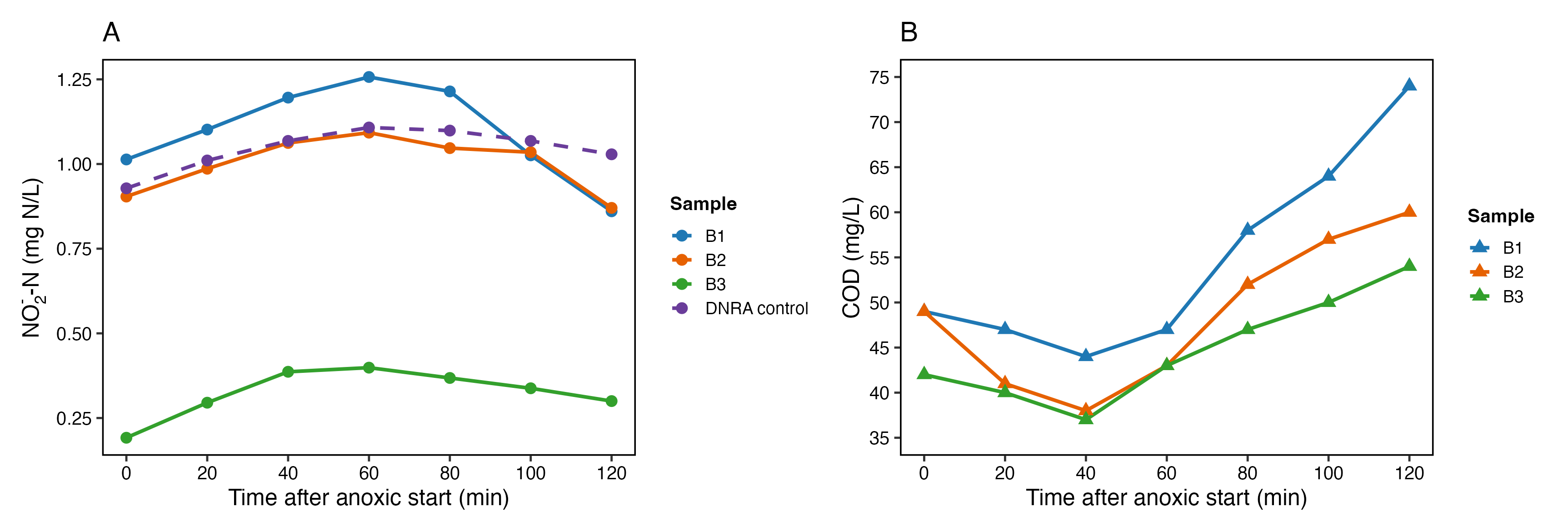

**Figure S4. (A) Nitrite dynamics during the anoxic stage in triplicate batch tests (B1–B3) and the DNRA control group. (B) COD dynamics during the anoxic stage in triplicate batch tests (B1–B3).**

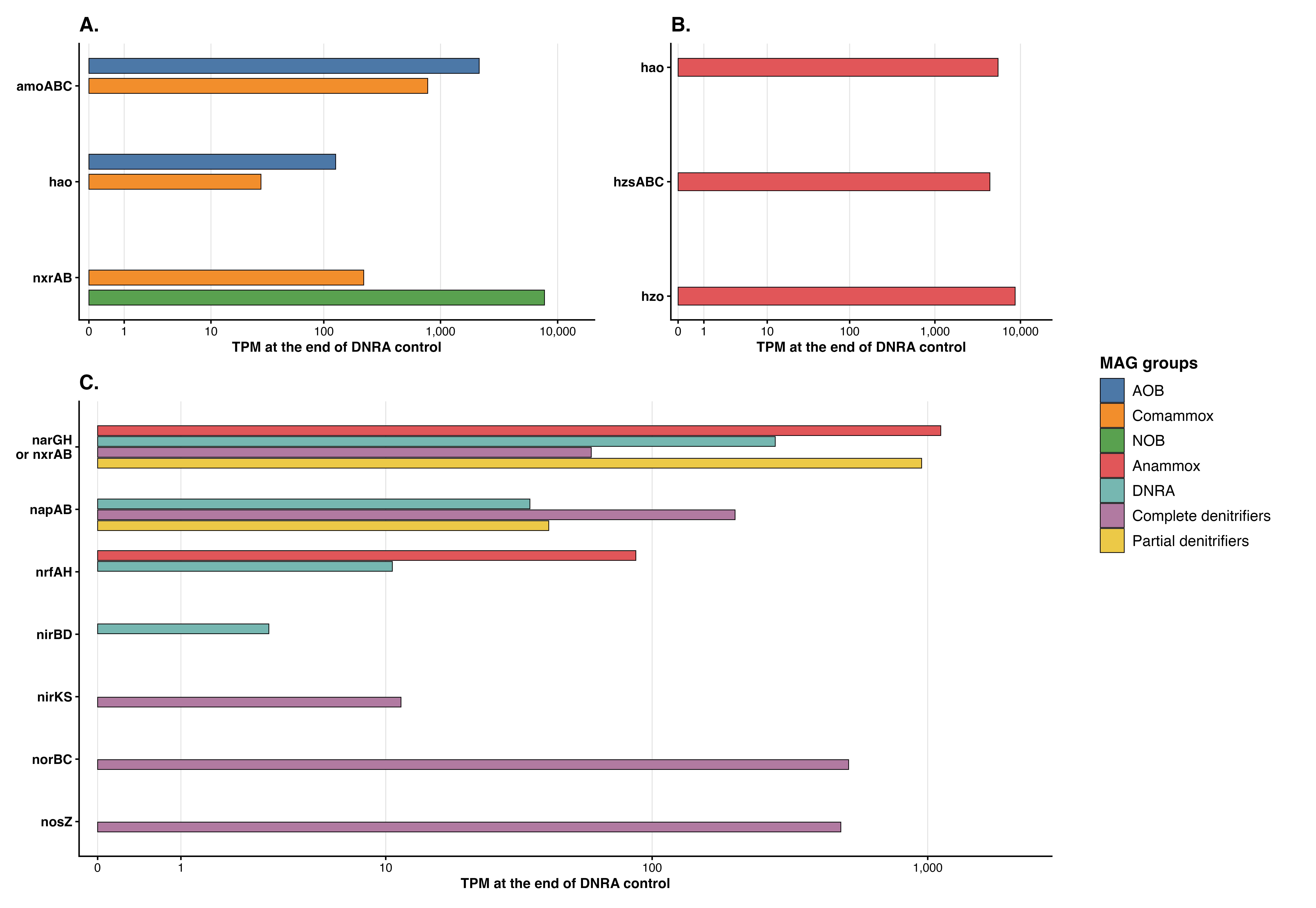

**Figure S5. Endpoint TPM values of nitrogen-metabolism genes affiliated with (A) nitrification-related bacteria, (B) anammox bacteria, and (C) nitrate- and nitrite-reducing MAG groups at the end of the representative DNRA control.**

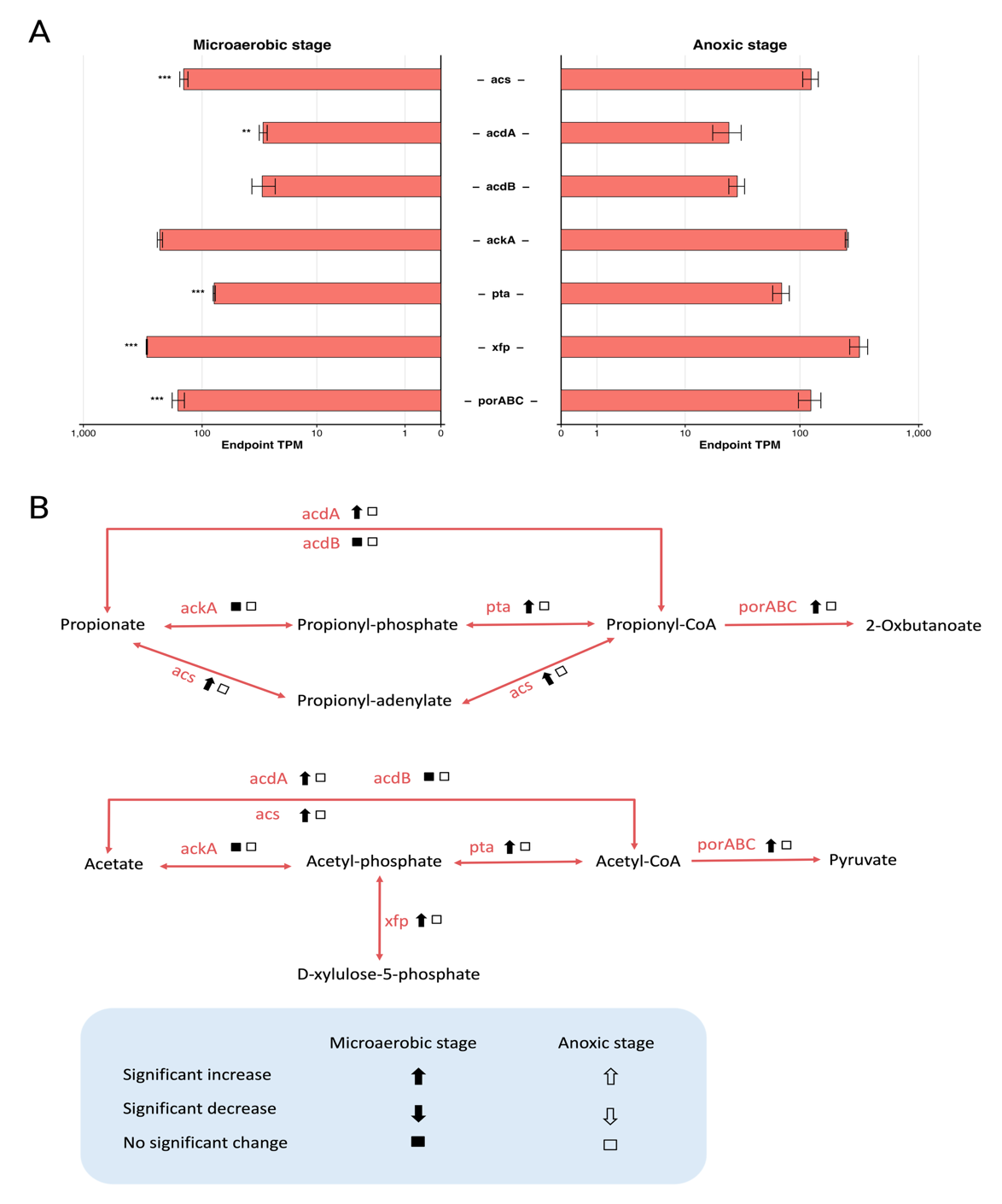

**Figure S6. (A) Endpoint TPM values of carbon-metabolism genes in the *Ca.* Brocadia sapporoensis MAG at the end of the microaerobic and acetate-amended anoxic stages.** Stars indicate significant transcriptional changes from the beginning to the end of each stage based on paired DESeq2 analysis of raw read counts after BH correction. **(B) Proposed acetate- and propionate-associated carbon-metabolism pathways inferred from the MAG annotation and transcriptional responses.** Solid arrows indicate significant increases, open arrows indicate nonsignificant increases, and open squares indicate no significant change.

**Table SI-1: Selected medium- and high-quality MAGs of nitrifiers, anammox bacteria, DNRA bacteria, and complete denitrifiers recovered from IFAS media collected from the aerobic zone.**

| MAG group | Genome | Completeness  (%) | Contamination  (%) | Length  (bp) | Contigs | GTDB Taxonomy |
| --- | --- | --- | --- | --- | --- | --- |
| AMX | SemiBin_8 | 100 | 0.58 | 3493538 | 1 | d__Bacteria;p__Planctomycetota;c__Brocadiia;o__Brocadiales; f__Brocadiaceae;g__Brocadia;s__Brocadia sapporoensis |
| AOB | 296 | 94.29 | 3.35 | 2995367 | 22 | d__Bacteria;p__Pseudomonadota;c__Gammaproteobacteria;o__Burkholderiales; f__Nitrosomonadaceae;g__Nitrosomonas;s__Nitrosomonas sp008015595 |
| CMX | SemiBin_67 | 90.91 | 8.42 | 4614235 | 63 | d__Bacteria;p__Nitrospirota;c__Nitrospiria;o__Nitrospirales; f__Nitrospiraceae;g__Nitrospira_D;s__Nitrospira_D sp002254365 |
| NOB | bin.536_sub | 88.45 | 0.7 | 3356908 | 5 | d__Bacteria;p__Nitrospirota;c__Nitrospiria;o__Nitrospirales; f__Nitrospiraceae;g__Nitrospira_D;s__Nitrospira_D sp900299245 |
| NOB | SemiBin_64 | 80.61 | 4.87 | 3297591 | 6 | d__Bacteria;p__Nitrospirota;c__Nitrospiria;o__Nitrospirales; f__Nitrospiraceae;g__Nitrospira_A;s__Nitrospira_A sp009594855 |
| NOB | SemiBin_37 | 89.44 | 0.94 | 3305409 | 10 | d__Bacteria;p__Nitrospirota;c__Nitrospiria;o__Nitrospirales; f__Nitrospiraceae;g__Nitrospira_D;s__ |
| NOB | SemiBin_2 | 99.93 | 0.61 | 3306502 | 1 | d__Bacteria;p__Nitrospirota;c__Nitrospiria;o__Nitrospirales; f__Nitrospiraceae;g__BQWY01;s__ |
| NOB | SemiBin_166 | 60.51 | 3.78 | 2719436 | 26 | d__Bacteria;p__Nitrospirota;c__Nitrospiria;o__Nitrospirales; f__Nitrospiraceae;g__Nitrospira_A;s__ |
| NOB | 930 | 69.51 | 0.19 | 1784591 | 22 | d__Bacteria;p__Pseudomonadota;c__Gammaproteobacteria; o__Burkholderiales;f__Gallionellaceae;g__Nitrotoga;s__ |
| NOB | 263 | 99.96 | 9.16 | 4585722 | 16 | d__Bacteria;p__Nitrospirota;c__Nitrospiria;o__Nitrospirales; f__Nitrospiraceae;g__Nitrospira_A;s__ |
| DNRA | 125 | 86.63 | 0.6 | 5873513 | 28 | d__Bacteria;p__Pseudomonadota;c__Gammaproteobacteria; o__Burkholderiales;f__Burkholderiaceae;g__Rubrivivax;s__ |
| DNRA | bin.664 | 72.92 | 1.81 | 2866143 | 61 | d__Bacteria;p__Acidobacteriota;c__Holophagae; o__Holophagales;f__Holophagaceae;g__Geothrix;s__Geothrix odensensis |
| DNRA | 926 | 75.28 | 1.22 | 4093228 | 44 | d__Bacteria;p__Pseudomonadota;c__Alphaproteobacteria; o__Rhizobiales;f__Hyphomicrobiaceae;g__;s__ |
| DNRA | 3 | 98.18 | 5.96 | 8349589 | 10 | d__Bacteria;p__Acidobacteriota;c__Terriglobia;o__Bryobacterales; f__Bryobacteraceae;g__JADLDI01;s__JADLDI01 sp020846745 |
| DNRA | 7 | 98.09 | 0.01 | 2722890 | 3 | d__Bacteria;p__Pseudomonadota;c__Gammaproteobacteria; o__Pseudomonadales;f__UBA5862;g__JACKOQ01;s__JACKOQ01 sp024234215 |
| DNRA | bin.222 | 76.65 | 3.43 | 2500900 | 45 | d__Bacteria;p__Pseudomonadota;c__Alphaproteobacteria; o__Sphingomonadales;f__Sphingomonadaceae;g__CANJQJ01;s__ |
| DNRA | bin.327 | 83.22 | 0.93 | 3699894 | 21 | d__Bacteria;p__Acidobacteriota;c__Vicinamibacteria; o__Vicinamibacterales;f__UBA2999;g__WHSN01;s__ |
| DNRA | bin.574 | 66.14 | 0.47 | 2523405 | 46 | d__Bacteria;p__Pseudomonadota;c__Alphaproteobacteria; o__Sphingomonadales;f__Sphingomonadaceae;g__;s__ |
| DNRA | bin.740 | 99.99 | 0.61 | 5549967 | 24 | d__Bacteria;p__Pseudomonadota;c__Alphaproteobacteria; o__Rhizobiales;f__Xanthobacteraceae;g__Bradyrhizobium;s__ |
| DNRA | bin.863 | 89.25 | 5.56 | 4337246 | 7 | d__Bacteria;p__Actinomycetota;c__Actinomycetes; o__Propionibacteriales;f__Propionibacteriaceae;g__Micropruina;s__ |
| DNRA | SemiBin_236 | 75.55 | 8.61 | 4549732 | 16 | d__Bacteria;p__Pseudomonadota;c__Gammaproteobacteria; o__Burkholderiales;f__Burkholderiaceae;g__Piscinibacter;s__ |
| DNRA | SemiBin_7 | 99.96 | 0.18 | 5606669 | 4 | d__Bacteria;p__Actinomycetota;c__Actinomycetes;o__Mycobacteriales; f__Mycobacteriaceae;g__Gordonia;s__Gordonia amarae |
| Complete denitrifiers | 134 | 87.34 | 2.21 | 3309181 | 14 | d__Bacteria;p__Pseudomonadota;c__Alphaproteobacteria;o__Rhizobiales; f__Hyphomicrobiaceae;g__Hyphomicrobium_B;s__Hyphomicrobium_B sp016793445 |
| Complete denitrifiers | 34 | 100 | 1.59 | 3790873 | 7 | d__Bacteria;p__Pseudomonadota;c__Alphaproteobacteria;o__Rhodobacterales; f__Rhodobacteraceae;g__Albidovulum;s__Albidovulum sp039808285 |
| Complete denitrifiers | bin.29 | 82.04 | 2.46 | 4853969 | 35 | d__Bacteria;p__Pseudomonadota;c__Gammaproteobacteria; o__Ga0077536;f__Ga0077536;g__Ga0077536;s__ |
| Complete denitrifiers | bin.618 | 91.03 | 2.74 | 3778957 | 40 | d__Bacteria;p__Pseudomonadota;c__Alphaproteobacteria; o__Rhodobacterales;f__Rhodobacteraceae;g__Albidovulum;s__Albidovulum sp030837825 |
| Complete denitrifiers | 17 | 98.82 | 3.07 | 3851201 | 5 | d__Bacteria;p__Pseudomonadota;c__Alphaproteobacteria; o__Rhizobiales;f__Hyphomicrobiaceae;g__Hyphomicrobium_B;s__ |
| Complete denitrifiers | 272 | 99.75 | 0.09 | 3127543 | 11 | d__Bacteria;p__Pseudomonadota;c__Gammaproteobacteria; o__Burkholderiales;f__Rhodocyclaceae;g__Azonexus;s__Azonexus sp016790845 |
| Complete denitrifiers | bin.420 | 99.95 | 1.58 | 3439739 | 9 | d__Bacteria;p__Pseudomonadota;c__Alphaproteobacteria;o__Rhizobiales; f__Hyphomicrobiaceae;g__Hyphomicrobium_B;s__Hyphomicrobium_B sp001006785 |

**Table S2. Table S2. Initial substrate concentrations and operational conditions for supplementary diagnostic control batch tests conducted without pH control.**

| Control Groups | NH_4_^+^  (mg N/L) | NO_2_^–^  (mg N/L) | NO_3_^–^  (mg N/L) | Acetate  (mg COD/L) | Continued Nitrogen Gas Sparging |
| --- | --- | --- | --- | --- | --- |
| A | 50 |  | 50 | 10 | Yes |
| B | 50 |  | 50 | 10 | No |
| C | 50 |  | 50 |  | Yes |
| D |  |  |  | 10 | Yes |
| E |  |  | 50 | 10 | Yes |
| F | 50 |  |  | 10 | Yes |

**Table S3. Nitrogen species, soluble COD, pH, temperature, and calculated free ammonia (FA) concentrations in supplementary diagnostic control batch tests conducted without pH control.**

| Control groups | NH4-N (mg/L) | | NO2-N (mg/L) | | NO3-N (mg/L) | | COD (mg/L) | | pH | | Temperature | Free Ammonia (mg/L) | |
| --- | --- | --- | --- | --- | --- | --- | --- | --- | --- | --- | --- | --- | --- |
|  | 0 min | 180 min | 0 min | 180 min | 0 min | 180 min | 0 min | 180 min | 0 min | 180 min | ℃ | 0 min | 180 min |
| A | 58.36 | 53.72 | 0.08 | 1.61 | 58.30 | 47.30 | 41.82 | 118.74 | 7.83 | 8.25 | 20.90 | 1.97 | 4.56 |
| B | 61.09 | 57.56 | 0.10 | 0.74 | 59.20 | 47.70 | 32.00 | 36.00 | 7.43 | 7.37 | 20.90 | 0.83 | 0.69 |
| C | 59.34 | 56.85 | 0.06 | 1.32 | 58.90 | 52.00 | 27.62 | 143.07 | 7.50 | 8.21 | 21.00 | 0.96 | 4.45 |
| D | 0.00 | 0.00 | 0.10 | 0.00 | 3.46 | 0.16 | 35.58 | 41.02 | 7.76 | 8.70 | 21.00 | 0.00 | 0.00 |
| E | 0.00 | 0.00 | 0.18 | 2.10 | 59.10 | 52.50 | 29.70 | 35.07 | 7.95 | 8.83 | 20.60 | 0.00 | 0.00 |
| F | 58.12 | 58.74 | 0.12 | 0.00 | 2.54 | 0.23 | 29.78 | 104.58 | 7.62 | 8.03 | 20.60 | 1.19 | 3.02 |

**Table S4. VFA concentrations in diagnostic control groups showing substantial pH or soluble COD increases under anoxic/anaerobic conditions.**

|  | A | | B | | C | | D | | E | | F | |
| --- | --- | --- | --- | --- | --- | --- | --- | --- | --- | --- | --- | --- |
| Time (min) | 0 | 180 | 0 | 180 | 0 | 180 | 0 | 180 | 0 | 180 | 0 | 180 |
| **Lactate (mg COD/L)** | ND | ND | ND | ND | ND | ND | ND | ND | ND | ND | ND | ND |
| **Formate (mg COD/L)** | ND | ND | ND | ND | ND | ND | ND | ND | ND | ND | ND | ND |
| **Acetate (mg COD/L)** | 5.12 | ND | 5.12 | ND | ND | ND | 8.32 | ND | 7.68 | ND | 8.32 | 8.96 |
| **Propionate (mg COD/L)** | 26.88 | ND | 10.08 | 15.68 | 26.88 | 17.92 | 17.92 | 13.44 | 8.96 | 11.20 | 16.80 | 13.44 |
| **Iso-butyrate (mg COD/L)** | ND | ND | ND | ND | ND | ND | ND | ND | ND | 35.20 | ND | ND |
| **Butyrate (mg COD/L)** | ND | 182.40 | ND | ND | ND | 107.20 | ND | 20.08 | ND | ND | ND | 176.00 |
| **Iso-valerate (mg COD/L)** | ND | ND | ND | ND | ND | ND | ND | ND | ND | ND | ND | ND |
| **Valerate (mg COD/L)** | ND | ND | ND | ND | ND | ND | ND | ND | ND | ND | ND | ND |
| **Methyl valerate (mg COD/L)** | ND | ND | ND | ND | ND | 17.92 | ND | ND | ND | ND | ND | ND |
| **Caproate (mg COD/L)** | ND | ND | ND | ND | ND | ND | ND | ND | ND | ND | ND | ND |

**Table S5. Initial substrate concentrations and operational conditions for supplementary free-ammonia control tests conducted under adjusted pH conditions.**

| Control Groups | Operation time (hour) | pH | NH_4_^+^  (mg N/L) | NO_3_^–^  (mg N/L) | Acetate  (mg COD/L) | Temperature ($℃$) | Theoretical FA Concentrations (mg/L) |
| --- | --- | --- | --- | --- | --- | --- | --- |
| 1 | 3 | 7 | 50 | 50 | 10 | 21.5 | 0.27 |
| 2 |  | 7.5 | 50 | 50 | 10 | 21.4 | 0.84 |
| 3 |  | 8 | 50 | 50 | 10 | 21.5 | 2.59 |
| 4 |  | 8.5 | 50 | 50 | 10 | 21.3 | 7.41 |
| 5 |  | 9 | 50 | 50 | 10 | 21.3 | 18.54 |

**Table S6. DESeq2-based differential expression and endpoint TPM values of nitrogen cycling-related gene modules during the microaerobic and acetate-amended anoxic stages.** Endpoint mean TPM values represent average transcript abundances at the end of each stage. Log2 fold changes were calculated relative to the corresponding stage startpoint using raw read counts, and adjusted p-values were obtained using the Benjamini–Hochberg correction. Modules not retained after low-count filtering are shown as NA.

| Genes | MAG groups | Stages | Endpoint mean TPM | DESeq2 log2FC | Adjusted p-value | Significant label |
| --- | --- | --- | --- | --- | --- | --- |
| amoABC | AOB | Microaerobic | 1815.80 | -0.19 | 1.99E-01 |  |
| hao | AOB | Microaerobic | 128.50 | -0.32 | 4.19E-01 |  |
| amoABC | CMX | Microaerobic | 585.96 | 0.02 | 9.53E-01 |  |
| hao | CMX | Microaerobic | 22.21 | 0.04 | 9.53E-01 |  |
| nxrAB | CMX | Microaerobic | 234.46 | 0.10 | 7.33E-01 |  |
| nxrAB | NOB | Microaerobic | 12221.41 | 0.36 | 5.04E-10 | *** |
| hao | AMX | Microaerobic | 5463.84 | 0.56 | 2.52E-15 | *** |
| hzsABC | AMX | Microaerobic | 4631.67 | -0.08 | 4.19E-01 |  |
| hzo | AMX | Microaerobic | 10515.83 | 0.55 | 5.01E-21 | *** |
| nxrAB | AMX | Microaerobic | 888.68 | 1.19 | 2.31E-21 | *** |
| nrfAH | AMX | Microaerobic | 86.92 | 0.68 | 2.66E-01 |  |
| narGH | Complete_DNRA_MAGs | Microaerobic | 153.08 | 1.14 | 5.87E-05 | *** |
| napAB | Complete_DNRA_MAGs | Microaerobic | 16.18 | 0.08 | 9.53E-01 |  |
| nrfAH | Complete_DNRA_MAGs | Microaerobic | 5.89 | 0.63 | 8.50E-01 |  |
| nirBD | Complete_DNRA_MAGs | Microaerobic | 2.11 | NA | NA | NA |
| narGH | Complete_Denitrifiers_MAGs | Microaerobic | 30.81 | 1.28 | 3.12E-02 | * |
| napAB | Complete_Denitrifiers_MAGs | Microaerobic | 19.21 | 0.79 | 4.53E-01 |  |
| nirKS | Complete_Denitrifiers_MAGs | Microaerobic | 2.78 | NA | NA | NA |
| norBC | Complete_Denitrifiers_MAGs | Microaerobic | 41.14 | 0.32 | 7.84E-01 |  |
| nosZ | Complete_Denitrifiers_MAGs | Microaerobic | 66.21 | 0.98 | 1.92E-02 | * |
| narGH | Partial_denitrification_MAGs | Microaerobic | 404.85 | 1.19 | 1.40E-11 | *** |
| napAB | Partial_denitrification_MAGs | Microaerobic | 16.26 | 0.24 | 8.50E-01 |  |
| amoABC | AOB | Anoxic | 1772.64 | 0.03 | 7.87E-01 |  |
| hao | AOB | Anoxic | 108.36 | -0.32 | 1.20E-01 |  |
| amoABC | CMX | Anoxic | 547.86 | -0.05 | 7.66E-01 |  |
| hao | CMX | Anoxic | 18.49 | -0.32 | 4.08E-01 |  |
| nxrAB | CMX | Anoxic | 205.90 | -0.24 | 1.20E-01 |  |
| nxrAB | NOB | Anoxic | 11513.42 | -0.13 | 3.24E-01 |  |
| hao | AMX | Anoxic | 6928.91 | 0.62 | 8.08E-06 | *** |
| hzsABC | AMX | Anoxic | 3501.90 | -0.20 | 2.39E-01 |  |
| hzo | AMX | Anoxic | 10876.19 | 0.31 | 1.47E-02 | * |
| nxrAB | AMX | Anoxic | 1760.25 | 1.38 | 2.47E-20 | *** |
| nrfAH | AMX | Anoxic | 99.43 | 0.55 | 3.05E-02 | * |
| narGH | Complete_DNRA_MAGs | Anoxic | 214.45 | 0.47 | 4.70E-03 | ** |
| napAB | Complete_DNRA_MAGs | Anoxic | 37.32 | 1.29 | 8.26E-05 | *** |
| nrfAH | Complete_DNRA_MAGs | Anoxic | 12.61 | 0.56 | 4.88E-01 |  |
| nirBD | Complete_DNRA_MAGs | Anoxic | 2.12 | NA | NA | NA |
| narGH | Complete_Denitrifiers_MAGs | Anoxic | 53.76 | 0.85 | 2.07E-04 | *** |
| napAB | Complete_Denitrifiers_MAGs | Anoxic | 113.58 | 2.36 | 3.27E-21 | *** |
| nirKS | Complete_Denitrifiers_MAGs | Anoxic | 14.14 | 1.81 | 8.70E-03 | ** |
| norBC | Complete_Denitrifiers_MAGs | Anoxic | 365.36 | 3.21 | 3.32E-64 | *** |
| nosZ | Complete_Denitrifiers_MAGs | Anoxic | 301.69 | 2.22 | 1.44E-46 | *** |
| narGH | Partial_denitrification_MAGs | Anoxic | 987.29 | 1.35 | 1.20E-25 | *** |
| napAB | Partial_denitrification_MAGs | Anoxic | 69.86 | 2.10 | 6.27E-16 | *** |

**Table S7. TPM-based transcriptional trends of nitrogen metabolism genes in the representative DNRA control.** Startpoint and endpoint TPM values were used to calculate descriptive TPM-based log2 fold changes. Because only one DNRA control was available for metatranscriptomic analysis, no statistical significance testing was performed.

| Genes | MAG Groups | Startpoint mean TPM | Endpoint mean TPM | TPM-based mean log2FC |
| --- | --- | --- | --- | --- |
| hao | AMX | 3670.75 | 5473.71 | 0.58 |
| hzo | AMX | 6639.39 | 8668.65 | 0.38 |
| hzsABC | AMX | 5294.15 | 4394.47 | -0.27 |
| nxrAB | AMX | 392.07 | 1114.73 | 1.51 |
| nrfAH | AMX | 40.44 | 87.19 | 1.09 |
| amoABC | AOB | 2497.25 | 2139.30 | -0.22 |
| hao | AOB | 263.19 | 126.54 | -1.05 |
| amoABC | CMX | 1024.37 | 777.38 | -0.40 |
| hao | CMX | 43.56 | 28.39 | -0.60 |
| nxrAB | CMX | 361.41 | 220.32 | -0.71 |
| napAB | Complete_DNRA_MAGs | 13.45 | 35.52 | 1.34 |
| narGH | Complete_DNRA_MAGs | 99.91 | 280.41 | 1.48 |
| nirBD | Complete_DNRA_MAGs | 3.40 | 3.16 | -0.08 |
| nrfAH | Complete_DNRA_MAGs | 1.70 | 10.62 | 2.11 |
| napAB | Complete_Denitrifiers_MAGs | 16.34 | 200.40 | 3.54 |
| narGH | Complete_Denitrifiers_MAGs | 12.74 | 59.85 | 2.15 |
| nirKS | Complete_Denitrifiers_MAGs | 2.06 | 11.49 | 2.03 |
| norBC | Complete_Denitrifiers_MAGs | 30.99 | 517.17 | 4.02 |
| nosZ | Complete_Denitrifiers_MAGs | 49.86 | 484.70 | 3.26 |
| nxrAB | NOB | 12619.38 | 7721.04 | -0.71 |
| napAB | Partial_denitrification_MAGs | 21.45 | 41.72 | 0.93 |
| narGH | Partial_denitrification_MAGs | 235.32 | 950.45 | 2.01 |

**Table S8. DESeq2-based differential expression and endpoint TPM values of carbon metabolism-related genes encoded by the *Ca.* Brocadia sapporoensis MAG during the microaerobic and acetate-amended anoxic stages.** Endpoint mean TPM values represent average transcript abundances at the end of each stage. Log2 fold changes were calculated relative to the corresponding stage startpoint using raw read counts, and adjusted p-values were obtained using the Benjamini–Hochberg correction.

| Genes | Stages | Endpoint mean TPM | DESeq2 log2FC | Adjusted p-value | Significant label |
| --- | --- | --- | --- | --- | --- |
| acs | Microaerobic | 143.03 | 0.94 | 1.36E-13 | *** |
| acdA | Microaerobic | 30.06 | 1.64 | 9.20E-03 | ** |
| acdB | Microaerobic | 30.55 | 0.33 | 4.60E-01 |  |
| ackA | Microaerobic | 227.27 | 0.21 | 7.57E-02 |  |
| pta | Microaerobic | 78.89 | 0.87 | 3.15E-09 | *** |
| xfp | Microaerobic | 292.66 | 0.46 | 1.71E-09 | *** |
| porABC | Microaerobic | 160.02 | 0.65 | 1.78E-06 | *** |
| acs | Anoxic | 124.29 | 0.13 | 7.70E-01 |  |
| acdA | Anoxic | 24.59 | -0.12 | 8.34E-01 |  |
| acdB | Anoxic | 29.09 | 0.30 | 7.70E-01 |  |
| ackA | Anoxic | 248.32 | 0.33 | 3.69E-01 |  |
| pta | Anoxic | 69.93 | -0.01 | 9.62E-01 |  |
| xfp | Anoxic | 317.92 | 0.31 | 3.69E-01 |  |
| porABC | Anoxic | 123.69 | -0.13 | 7.70E-01 |  |

**Reference:**

Aktan, C.K., Yapsakli, K., Mertoglu, B., 2012. Inhibitory effects of free ammonia on Anammox bacteria. Biodegradation 23, 751–762. https://doi.org/10.1007/s10532-012-9550-0

Anthonisen, A., Loehr, R., Prakasam, T., Srinath, E., 1976. Inhibition of nitrification by ammonia and nitrous acid. Journal (Water Pollution Control Federation) 835–852.

Cotto, I., Vilardi, K.J., Huo, L., Fogarty, E.C., Khunjar, W., Wilson, C., De Clippeleir, H., Gilmore, K., Bailey, E., Lücker, S., Pinto, A.J., 2023. Low diversity and microdiversity of comammox bacteria in wastewater systems suggest specific adaptations within the Ca. Nitrospira nitrosa cluster. Water Research 229, 119497. https://doi.org/10.1016/j.watres.2022.119497

Hejnfelt, A., Angelidaki, I., 2009. Anaerobic digestion of slaughterhouse by-products. Biomass and Bioenergy 33, 1046–1054. https://doi.org/10.1016/j.biombioe.2009.03.004

Jaroszynski, L.W., Cicek, N., Sparling, R., Oleszkiewicz, J.A., 2012. Impact of free ammonia on anammox rates (anoxic ammonium oxidation) in a moving bed biofilm reactor. Chemosphere 88, 188–195. https://doi.org/10.1016/j.chemosphere.2012.02.085

Kayhanian, M., 1999. Ammonia Inhibition in High-Solids Biogasification: An Overview and Practical Solutions. Environmental Technology 20, 355–365. https://doi.org/10.1080/09593332008616828

Liu, Y., Ngo, H.H., Guo, W., Peng, L., Wang, D., Ni, B., 2019. The roles of free ammonia (FA) in biological wastewater treatment processes: A review. Environment International 123, 10–19. https://doi.org/10.1016/j.envint.2018.11.039

Yang, G., Xu, Q., Wang, D., Tang, L., Xia, J., Wang, Q., Zeng, G., Yang, Q., Li, X., 2018. Free ammonia-based sludge treatment reduces sludge production in the wastewater treatment process. Chemosphere 205, 484–492. https://doi.org/10.1016/j.chemosphere.2018.04.140

Zhang, C., Qin, Y., Xu, Q., Liu, X., Liu, Y., Ni, B.-J., Yang, Q., Wang, D., Li, X., Wang, Q., 2018. Free Ammonia-Based Pretreatment Promotes Short-Chain Fatty Acid Production from Waste Activated Sludge. ACS Sustainable Chem. Eng. 6, 9120–9129. https://doi.org/10.1021/acssuschemeng.8b01452

Zhao, J., Liu, Y., Wang, Y., Lian, Y., Wang, Q., Yang, Q., Wang, D., Xie, G.-J., Zeng, G., Sun, Y., Li, X., Ni, B.-J., 2018. Clarifying the Role of Free Ammonia in the Production of Short-Chain Fatty Acids from Waste Activated Sludge Anaerobic Fermentation. ACS Sustainable Chem. Eng. 6, 14104–14113. https://doi.org/10.1021/acssuschemeng.8b02670
